# Nanopore tweezers measurements of RecQ conformational changes reveal the energy landscape of helicase motion

**DOI:** 10.1101/2021.09.23.461529

**Authors:** Jonathan M. Craig, Maria Mills, Andrew H. Laszlo, Hwanhee C. Kim, Jesse R. Huang, Sarah J. Abell, Jonathan W Mount, Keir C. Neuman, Jens H. Gundlach

## Abstract

Helicases are essential for nearly all nucleic acid processes across the tree of life. Using Nanopore Tweezers we observed the small, fast steps taken by single RecQ helicases as they step along and unwind DNA at ultrahigh spatiotemporal resolution. By directly measuring conformational substates of RecQ we determine the coupling between helicase domain motions and chemical reactions that together produce forward motion along the DNA. Application of assisting and opposing forces shows that RecQ has a highly asymmetric energy landscape that reduces its sensitivity to opposing mechanical forces that could be encountered *in vivo* by molecular roadblocks such as DNA bound proteins. This energy landscape enables RecQ to maintain speed against an opposing load.

## Main Text

Helicases are molecular machines that hydrolyze ATP to unwind or remodel nucleic acids. They are involved in all aspects of genome maintenance, from replication and transcription to damage repair and removing bound proteins (*1–3*). The highly conserved family of RecQ helicases are essential for genome integrity. In humans, mutations in RecQ helicases are associated with several disorders characterized by premature aging and a predisposition to cancer (*4*). RecQ helicases, of which *E. coli* RecQ is the archetype (*3, 5–9*), unwind double-stranded DNA (dsDNA) by translocating 3′ to 5′ on one strand while displacing the other strand (*4, 10, 11*). RecQ helicases belong to the Super Family 2 (SF2) class of helicases, which use two highly conserved RecA-like domains to couple ATP binding and hydrolysis to translocation in an inchworm-like fashion (*12, 13*).

Single-molecule experiments have revealed molecular details of how RecQ helicases use the energy of ATP hydrolysis to generate mechanical work (*14–17*). However, because *E. Coli* RecQ moves in fast, small steps (~100 nt/s with ~1 nt step size), resolving individual steps of RecQ is beyond the spatial and temporal resolution of most single-molecule approaches. Here we leverage the spatiotemporal resolution of Single-molecule Picometer Resolution Nanopore Tweezers (SPRNT) (*18–20*) to characterize the kinetics and mechanochemistry of *E. Coli* RecQ helicase at previously unresolvable detail. In SPRNT, a single MspA nanopore is used to monitor the progression of a motor enzyme moving on DNA (Fig. 1, materials and methods, Fig. S1, Table S1-S2). The voltage applied across the pore establishes an electric field within the pore constriction that causes a sequence-dependent ion current to flow through the pore. The ion current is then converted to the nucleotide position of the motor enzyme (*18, 19*). The electric field applies a force to negatively charged DNA that is proportional to the applied voltage. This force can either assist or oppose the motion of the motor depending on whether the activity of the helicase results in DNA motion in the direction of, or opposite of, the electric force. In contrast to other force-based techniques such as optical or magnetic tweezers, which typically bias motor motion indirectly by destabilizing the DNA duplex, in SPRNT an assisting or hindering force is applied directly to the motor as it moves along the DNA. Furthermore, SPRNT is uniquely able to measure internal domain motions of the motor that alter the position of the DNA within the pore (*20*).

**Figure 1:**
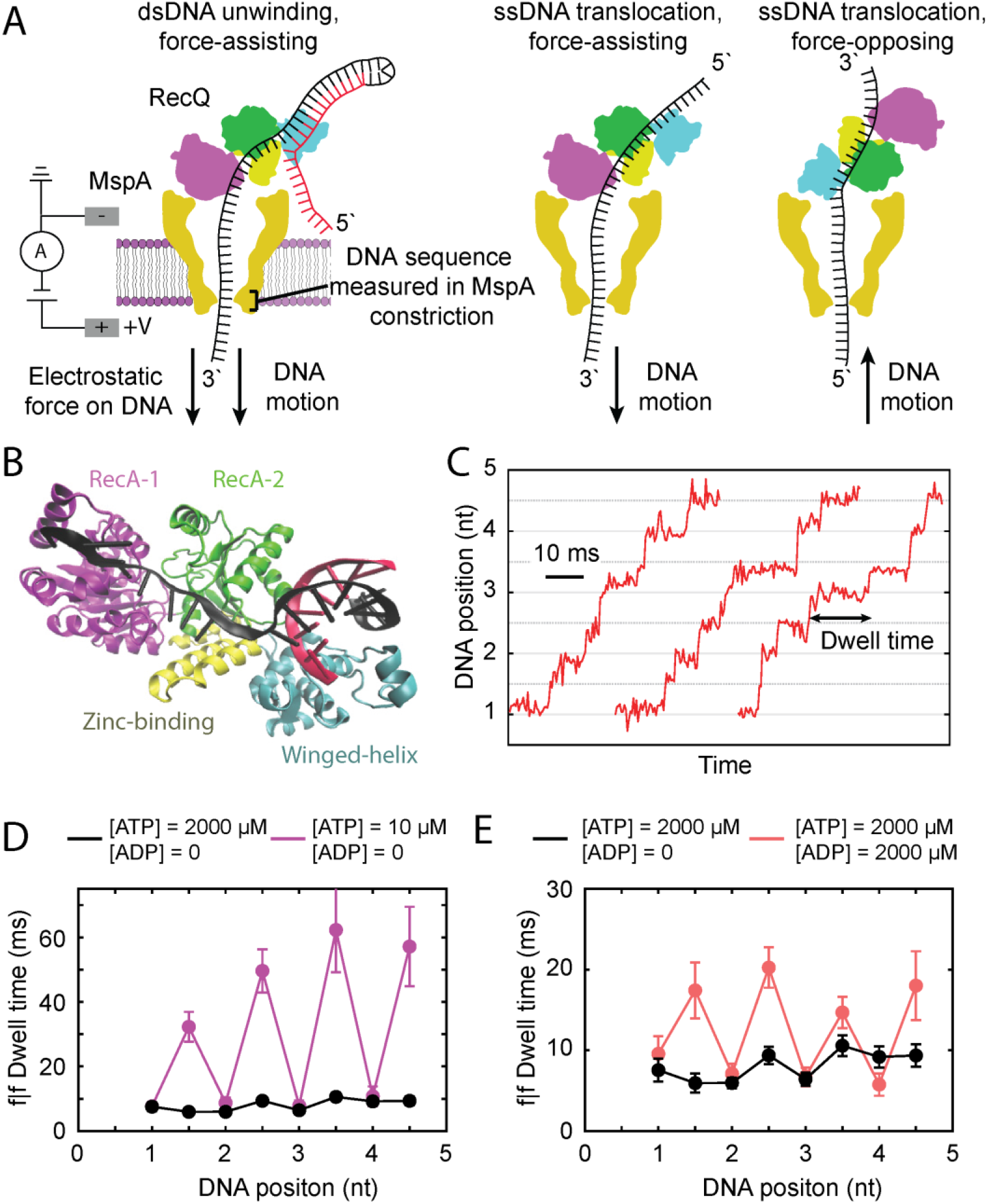
SPRNT measurements of RecQ helicase. (A) Left, a single RecQ unwinding dsDNA is drawn into the MspA pore via the single-stranded 3’ end of the template. The force on the DNA is in the direction of the DNA motion, leading to an assisting force. Middle, RecQ translocation on ssDNA with the 3’ end through the pore. Right, RecQ translocation on ssDNA with the 5’ end through the pore. (B) Crystal structure of RecQ from *C. Sakazakii* bound to dsDNA (PDB 6CRM). Protein domains are: RecA1 (pink), RecA2 (green), Zinc binding (yellow), and Winged helix (blue). (C) SPRNT position vs. time traces for three RecQ unwinding events. (D) Dwell-time vs. DNA position for the DNA positions shown in Figure 1C at [ATP] = 2000 µM (black) and [ATP] = 10 µM (pink). (E) Dwell-time vs. DNA position at [ATP] = 2000 µM with [ADP] = 0 (black) and at [ATP] = 2000 µM with [ADP] = 2000 µM (red). All errors are SEM.

We used SPRNT previously to determine the kinetic mechanism of ssDNA translocation of another Superfamily 2 helicase, Hel308, against an opposing force (*20, 21*). We observed two distinct kinetic sub-steps per nucleotide translocated by Hel308. One sub-step was ATP-concentration-dependent ([ATP]-dependent) whereas the second was ATP-concentration-independent ([ATP]-independent). We hypothesized these two states corresponded to open and closed conformations of the Hel308 RecA-like domains, respectively. Here, we expand and extend these results using RecQ. We directly observe and quantify the coupling between the ATP hydrolysis cycle and RecQ domain motion, and we use force spectroscopy to map the full energetic landscape of RecQ translocation. These results generalize our previous results obtained with Hel308 and suggest that SF2 helicases employ a common translocation mechanism.

We measured RecQ motion on DNA in three distinct geometries (Fig. 1A): (1) Force-assisted unwinding of a DNA hairpin, (2) force-assisted translocation on ssDNA and, (3) force-opposed translocation on ssDNA (Fig S2). Kinetic analysis of this data reveals how RecQ couples ATP binding and hydrolysis to internal domain motions that power DNA translocation and provides the most detailed picture to date of the energetic landscape of a helicase proceeding through its coupled ATP hydrolysis and translocation pathways

In force-assisted unwinding experiments, we observed two approximately half-nucleotide steps per nucleotide translocated through the nanopore by RecQ (Fig. 1C). We used ATP and ADP titration experiments to correlate these kinetic sub-steps in the ATP hydrolysis cycle with the RecA-like domain motions that are directly resolved in the SPRNT traces (Fig. 1D/E). Specifically, we measured the ATP and ADP concentration dependence of dwell times at integer and half-integer steps for a forward step following a forward step (f|f), which corresponds to processive stepping of the helicase (*20, 22, 23*). We found that the half-integer states are [ATP]-dependent whereas the integer steps are [ATP]-independent (Fig. 1D). In ADP titration experiments, ADP acts as a competitive inhibitor to progression of the [ATP]-dependent state (Fig. 1E). This suggests that ADP unbinding/binding occurs during the [ATP]-dependent state, during which it competes with ATP binding (Fig S3).

These results can be interpreted within the context of the inchworm model of SF2 helicase motion (*24, 1, 6*). In this model, ATP binding in the [ATP]-dependent (open) state drives a conformational change in the RecA-like domains from open to closed. ATP is hydrolyzed in the [ATP]-independent (closed) state, after which the RecA-like domains return to the open conformation, followed by ADP release, completing one catalytic cycle. Coordinated changes in the relative ssDNA binding energies of the RecA-like domains in response to these chemical reactions leads to directed motion along the DNA. The similarity of these results to those obtained with Hel308 suggest this may be a universal feature of processive SF2 helicase motion.

Additional insights into the RecQ hydrolysis cycle were obtained using a slowly-hydrolyzable ATP analogue, ATPγS, which induces ~1 second-long pauses in RecQ translocation (*17*). In unwinding experiments with a 2:1 ratio of ATP to ATPγS, we observed pauses of ~50 ms to > 1 s (Fig. 2A). Two distinct pause behaviors were associated with ATPγS binding depending on the position of RecQ on the DNA. We found that at nucleotide positions 1 and 4, RecQ exhibited long pauses (~1 s) in the [ATP]-independent state, followed by a forward transition to the next [ATP]-dependent state (Fig 2A). At nucleotide positions 2 and 3, RecQ exhibited shorter pauses (~150 ms) in the [ATP]-independent state with transient backwards steps to the previous [ATP]-dependent state, before eventually moving forwards via a short dwell-time state (Figs. 2B, 2C). These final, short dwell-time steps following a pause had similar dwell-times to those observed in the absence of ATPγS (Fig 2A-2C). This position dependence of the ATPγS-induced pausing behaviors suggests that RecQ kinetics are sensitive to the DNA sequence (Fig S4).

**Figure 2:**
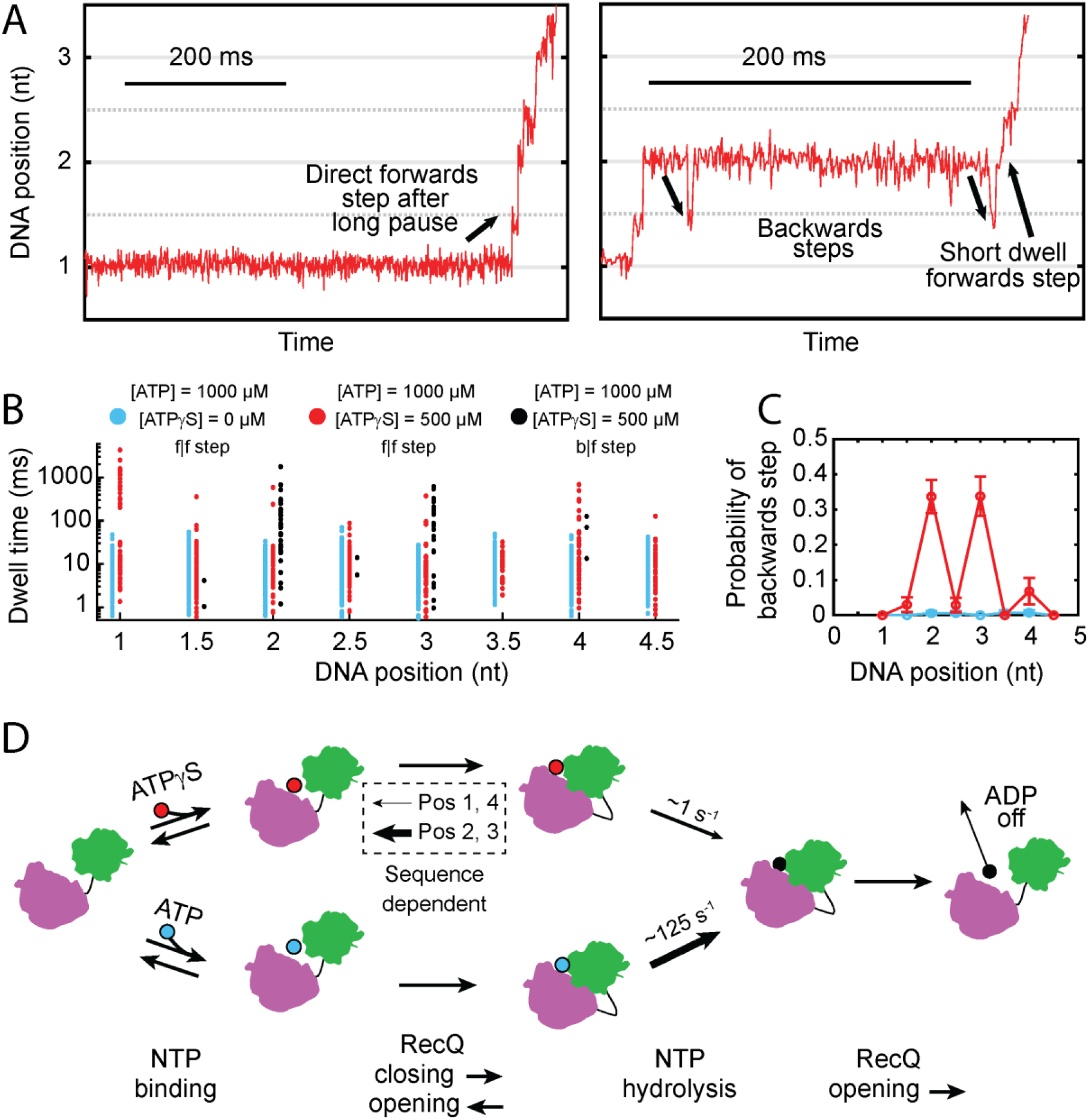
Sequence-dependent RecQ kinetics revealed by ATPγS binding. (A) Position vs. time traces demonstrating two different ATPγS-induced RecQ pause behaviors. (B) Dwell-times vs. DNA position for f|f steps at [ATP] = 1000 µM and [ATPγS] = 0 µM (blue), f|f steps in the presence of [ATPγS] = 500 µM (red) and b|f steps in the presence of ATPγS (black). (C) Probability of backwards step vs. DNA position at [ATP] = 1000 µM and [ATPγS] = 0 µM (blue) and at [ATP] = 1000 µM and [ATPγS] = 500 µM (red). (D) RecQ ATPγS hydrolysis pathway. The RecA-like domains of RecQ are represented in purple and green. ATP and ATPγS compete for the binding site at the interface of the RecA-like domains. Upon NTP binding the RecA-like domains adopt a closed conformation. If ATP is bound, RecQ continues through the hydrolysis pathway. If ATPγS is bound, RecQ can either slowly hydrolyze ATPγS (positions 1,4) or transiently open and eject ATPγS (positions 2,3).

These observations lead to a refined model in which ATPγS binds to RecQ in the [ATP]-dependent state, triggering a conformational change that closes the RecA-like domains. Depending on the position of RecQ on the nucleotide sequence, the helicase either slowly hydrolyzes ATPγS (Fig. 2D, positions 1 and 4) or alternates between conformational states (opening and closing of the RecA-like domains), eventually ejecting the ATPγS and allowing an ATP molecule to bind before proceeding through the remainder of the ATP hydrolysis cycle (Fig. 2D, positions 2 and 3). The data lend further support to our inference that ATP hydrolysis occurs during the [ATP]-independent state (*20*). The reported ATPγS hydrolysis rate by RecQ is ~0.2 s^−1^, though the ATPγS off rate is much faster and is the primary pause escape pathway (*17*). Our results are consistent with previous measurements but provide additional insights. RecQ adopts a closed conformation during ATPγS-induced pausing; the DNA sequence modifies the pause behavior by shifting the equilibrium between open and closed conformations; and opening of the RecA-like domains is the rate-limiting step in ejecting ATPγS.

The position dependent dwell-time data allowed us to construct a complete kinetic model of the RecQ mechanochemical cycle (Fig. 3A, supplementary text, Fig S5-S9 Table S3). We estimated each rate-constant in our model by likelihood maximization (Fig S10-S13, Table S4-S5). Comparison of our RecQ model with that derived previously for Hel308 suggests the mechanism of translocation applies broadly to processive SF2 helicases, but also reveals differences in off-pathway behaviors. Moreover, the RecQ results directly link the [ATP]-independent state with ATP hydrolysis, which could only be conjectured for Hel308 (*20*).

**Figure 3:**
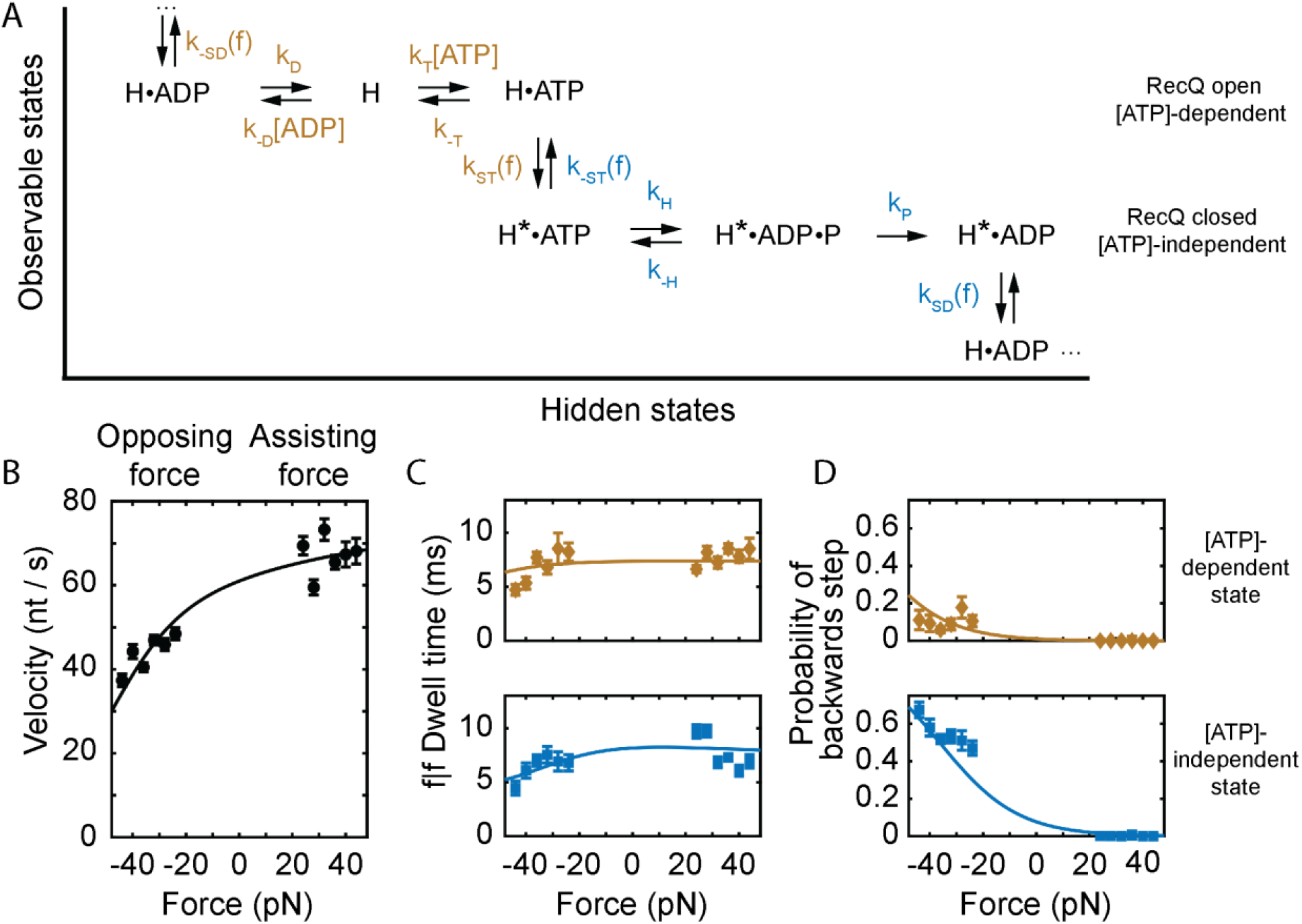
Force-dependent kinetics of RecQ translocation on ssDNA at saturating [ATP]. (A) Kinetic model of RecQ translocation on ssDNA. Rate-constants in gold affect the kinetics of the [ATP]-dependent step, rate-constants in blue affect the [ATP]-independent step. ‘H’ represents RecQ in the open conformation whereas ‘H*’ indicates RecQ in the closed conformation. (B) RecQ velocity vs. force. (C) f|f dwell time vs. force for the [ATP]-dependent state (gold, top) and the [ATP]-independent state (blue, bottom). (D) Probability of backwards step vs. force for the [ATP]-dependent state (gold, top) and the [ATP]-independent state (blue, bottom). Solid lines in (A, B, C) are global fits to our kinetic model (Fig. 3D). All errors are SEM.

In addition to the details of the kinetic pathway revealed by the unwinding experiments, our ability to apply both assisting and opposing force enables determination of the location and size of the energetic barriers between states (Fig S8). We measured RecQ translocation on ssDNA in force-assisting and force-opposing configurations to understand how RecQ generates and responds to force and to determine the energy landscape of RecQ motion. We used a poly-T ssDNA template to minimize sequence-dependent effects (Fig S2).

The ssDNA and dsDNA translocation rates were similar, consistent with previous descriptions of RecQ as an active helicase (*25*). The translocation velocity of RecQ remained constant with assisting force but decreased with opposing force (Fig 3B). Surprisingly, the decrease in velocity is not the result of a decrease in the forwards stepping rate, but rather due to an increase in the probability of transient backwards steps. The f|f dwell-times of both [ATP]-dependent and [ATP]-independent sub-states are weakly dependent on force (Fig 3C), whereas the probability of a backwards step is ~0 at high assisting forces, but increases dramatically at high opposing forces, particularly in the [ATP]-independent state (Fig 3D).

These results can be interpreted in the context of transition state theory in which the coupling of kinetics to force depends on *both* the magnitude of the applied force (voltage) and the distance to the transition state (supplementary text). There are two mechanical transitions associated with RecQ conformational changes that perform work by moving DNA through the pore: opening and closing of the RecA-like domains with ATP bound and ADP bound. The corresponding rate constants are therefore sensitive to the applied force. The backwards rates (k_-SD_ and k_-ST_) are far more sensitive to applied force than the forwards rates (Fig. 4A), suggesting an asymmetric energy landscape in which both transition states are located much closer to the pre-translocated state than to the post-translocated state (Figure 4B). This has a profound effect on the response of the helicase to mechanical force, dramatically reducing the sensitivity of the velocity to opposing forces, suggesting that RecQ may be optimized to work against opposing loads (Figure 4C).

**Figure 4:**
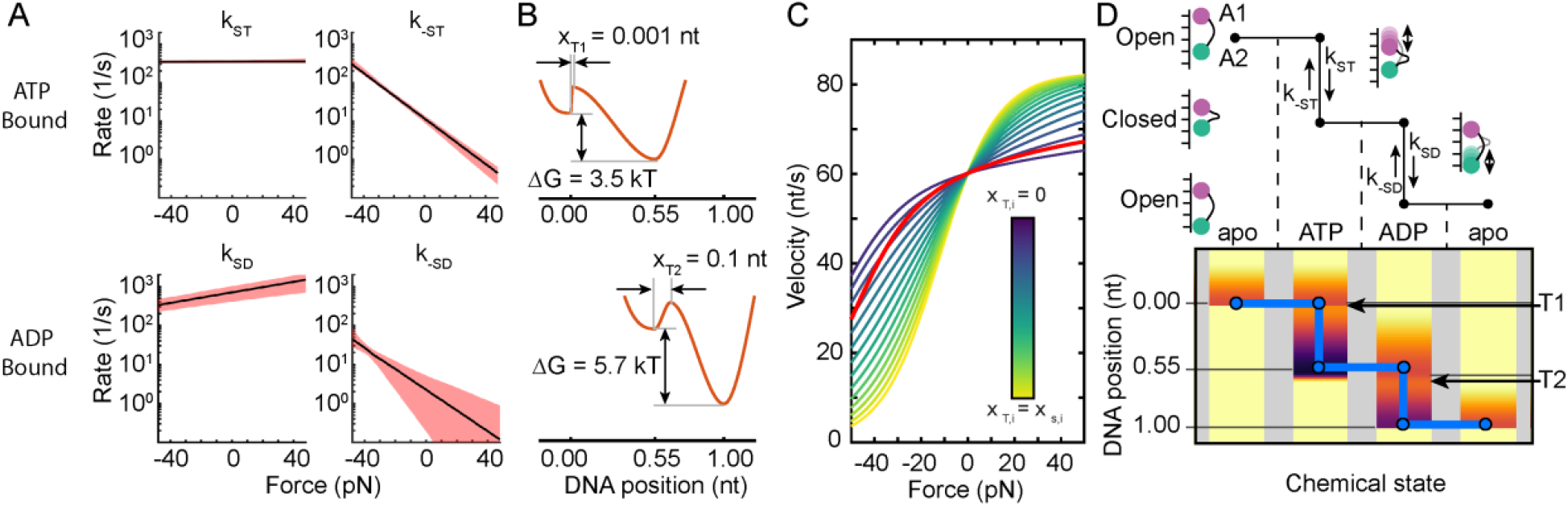
The energy landscape of DNA translocation by RecQ helicase. (A) Force-dependence of rate constants corresponding to conformational changes of (Top) ATP-bound RecQ and (Bottom) ADP-bound RecQ. Shaded regions are 95% confidence intervals. (B) Energy landscape diagram derived from the kinetic parameters in A. (C) RecQ velocity vs. force as a function of covaried transition state. Blue curves correspond to both transition states located at the pre-translocated state, and yellow curves to both transition states located at the post-translocated state. The red line is derived from our data. (D) Combined kinetic model for RecQ motion along ssDNA showing how the enzyme conformation and nucleotide bound state relate to the energy landscapes in (B) depicted in shaded colors below. Darker colors represent lower energy. T1 and T2 correspond to the location of the transition states from the ATP and ADP bound states, respectively.

From these data we constructed a free energy diagram for the RecQ hydrolysis cycle. The reaction coordinate is the DNA position in the MspA nanopore, with the measured position states corresponding to energy minima. The transition state positions and energy differences between the minima are determined by fitting the kinetic model to the data (Fig. 4A-C supplementary text). Figure 4D shows how the different energy landscapes associated with different nucleotide bound states relate to RecQ motion. In the apo state (no bound ATP or ADP), RecQ is in an open conformation and no steps occur. ATP binding alters the energy landscape (Fig. 4B), making a forward step favorable. RecQ then adopts a closed configuration that results in DNA advancing through the pore. The closed conformation enables ATP hydrolysis, again changing the energy landscape (Fig. 4D). This change disallows backwards steps, instead favoring another forward step associated with opening of the RecA-like domains. Once open, ADP release completes the hydrolysis cycle and RecQ returns to the apo state, generating a net motion of one full nucleotide along the DNA.

These results reveal how RecQ couples ATP binding and hydrolysis to DNA translocation in unprecedented detail. Whereas the mechanism we observe is consistent with structural and biochemical findings, it was not previously accessible experimentally. With SPRNT, we follow both the RecA-like domain motions and DNA translocation, which enables the mechanochemical coupling between them to be determined directly with high spatiotemporal resolution. By determining the effects of force on the kinetics of mechanical steps, we mapped the entire energy landscape. The unusual energy landscape suggests that RecQ is optimized to work against opposing force (Fig S14), for example in stripping bound proteins from ssDNA. The energy landscape also explains the insensitivity of RecQ’s velocity to assisting forces (Fig. 3,4).

The core mechanism of RecQ translocation appears to apply generally to processive SF2 helicases. Nonetheless, important mechanistic differences underlying their different physiological roles exist among these helicases which can be resolved using SPRNT analysis.

Single molecule experiments provide maximum detail of the mechanical and chemical activities of enzymes. This work demonstrates the ability of SPRNT to resolve details of chemical and physical pathways of DNA translocating enzymes beyond what was previously accessible. The SPRNT experimental and analytical framework developed here to interpret RecQ behavior can be broadly applied to infer mechanism and quantify kinetic and energetic parameters in other enzyme systems.

## Acknowledgements

We thank Gabor Harami & Mihaly Kovacs for providing some of the RecQ enzyme used in this study.

## Supplementary Materials

Materials and Methods

Table S1-S5

Fig S1-S14 References (*15, 18–23, 26–28*)

## Author contributions

JMC – conceptualization, methodology, software, formal analysis, data collection, writing, visualization, supervision

MM – conceptualization, methodology, writing, visualization, supervision, resources

AHL – software, formal analysis, writing, visualization, supervision

SJA – data collection

HCK – data collection

JRH – data collection

JWM – data collection

KCN – conceptualization, methodology, writing, visualization, supervision, resources

JHG – funding acquisition

