## Supplementary Text for "Nanopore tweezers measurements of RecQ conformational changes reveal the energy landscape of helicase motion"

Contents


1. Materials and Methods 2
1.1. Pore Establishment 2
1.2. Operating Conditions 2
1.3. Proteins 2
1.4. DNA preparation 2
1.5. Data Acquisition 2
1.6. Raw Data Analysis 2
1.7. Kinetic Methods 2
1.8. Accounting for short-duration state loss in dwell time calculations 2
1.9. DNA design 5
1.10. Experimental Statistics 7
2. Competitive inhibition model for [ATP]-dependent state 8
3. Analysis of sequence-dependent effects on ATP$\gamma$S hydrolysis 9
4. Identification of [ATP]-dependent states in force-opposing configuration 12
5. Mechanism of $b|f$ [ATP]-independent steps; ruling out a futile hydrolysis pathway 14
6. Step size measurements in force-assisting and force-opposing ssDNA translocation experiments 15
7. Modelling force-dependent rate constants in SPRNT; a constraint equation 16
8. Parameter estimation in force-dependent RecQ translocation on ssDNA 18
9. Estimate of $k_{-T}$ and $k_T$ from ATPase data 24
10. Calculating the coordinates of the transition states and $\Delta G$ from kinetic parameters. Constructing the Energy Diagram 26
11. Exploring how the transition state coordinate affects RecQ kinetics 27

### 1. Materials and Methods

1.1. **Pore Establishment.** A single M2-NNN MspA nanopore was established in a 1,2-di-O-phytanyl-sn-glycero-3-phosphocholine (DOPHPC) lipid bilayer using methods that have been well established (Laszlo *et al.* 2016 Methods). Lipids were from Avanti Polar Lipids.

1.2. **Operating Conditions.** All experiments were run at 500 mM *trans* [KCl] and 100 mM *cis* [KCl] with 10 mM HEPES at pH 8.0 and 10 mM [$MgCl_2$] (*cis* only) at room temperature $22 \pm 1$ degrees Celsius. Once a single M2-NNN MspA nanopore was established, a buffer with the above conditions along with ATP, ADP and ATP$\gamma$S was perfused to the *cis* well. ATP, ADP and ATP$\gamma$S were ordered from Sigma Aldrich. The perfusion is done to maintain constant concentrations of the reactants/products in the reaction volume. DNA, DTT, and RecQ were added to final concentrations of 10 nM, 1 mM, and 50 nM, respectively. Reactants/products were re-perfused every 45 minutes.

1.3. **Proteins.** M2-NNN MspA (accession number CAB56052.1) were prepared as described previously (Derrington *et al.* 2015 Nature Biotechnology, Laszlo *et al.* 2016 Methods). We used a RecQ mutant lacking the Helicase-and-RNaseD-C-terminal (HRDC), whose preparation is described in (Harami *et al.* 2017 PNAS).

1.4. **DNA preparation.** Full DNA design concepts are discussed in section 1.9. DNA strands were suspended in 100 mM KCl at 20 $\mu M$ concentration. Hairpin DNA sequences were diluted further to 1 $\mu M$ and annealed by heating to 90°C and cooled to 4°C over 4 minutes. Sequences which required binding to a complement were annealed at 90°C and then decreased step-wise by $\approx$ 10°C per minute to 4°C. Once annealed these sequences were diluted to 1 $\mu M$ for addition to the experiment.

1.5. **Data Acquisition.** Data was acquired with custom labview software on an Axopatch 200B amplifier at 50 kHz, and downsampled by averaging to 25 kHz or 10 kHz.

1.6. **Raw Data Analysis.** Ion-current vs. time raw traces were converted to position vs. time traces as described in (Derrington *et al.* 2015, Laszlo *et al.* 2016 Methods, Craig *et al.* 2017 PNAS). At saturating [ATP] typical events were observed on the nanopore for several hundred milliseconds and $\approx$ 10 nt distance traveled. dwell times were determined by aligning ion-current segments to a consensus sequence as in (Craig *et al.* 2017 PNAS, Craig *et al.* 2019 Nucleic Acids Research).

1.7. **Kinetic Methods.** Previous work has shown that the observable dwell times depend on the chemical path leading into and out of an observable state (Tsygankov *et al.* 2007 Phys. Rev. E, Chemla *et al.* 2008 J. Phys. Chem. B, Craig *et al.* 2017 PNAS, Craig *et al.* 2021 Essays in Biochemistry). We use the following notation to denote enzyme step types: $f|f$ (forwards step following forwards step), $f|b$ (forwards step following backwards step) and $b|f$ (backwards step following forwards step). Observable states in SPRNT records are typically composed of several chemical or physical subprocesses, such as ATP binding and a conformational change. A $f|f$ step is the enzyme's typical processive step and proceeds through several subprocesses, while an $f|b$ step may follow only a subset of these processes, altering the dwell time distribution. By analyzing how conditional step types respond to experimental conditions, we make inferences about an enzyme's kinetic pathways.

1.8. **Accounting for short-duration state loss in dwell time calculations.** The measurements of half-nucleotide steps at millisecond duration approach SPRNT's spatio-temporal resolution. As a result, ion-current states ('levels') can be missed by level-finding algorithms. Here we estimate the time-scale at which a significant fraction of the levels are missed, and develop a method for correcting the mean dwell time that accounts for these missed short-dwell time states. As an example, we consider the case of f|b [ATP]-dependent states. We have found that the distribution of f|b steps is skewed towards short dwell times compared to f|f steps (Fig s6). We model the distribution of f|b steps as a a single exponential distribution and look for deviations from that model to estimate the fraction of states lost as a function of dwell time. The count-normalized distribution function takes the form

$$\rho_{f|b} = N_0 \cdot exp(t/\langle t \rangle) \tag{s1}$$

We binned the dwell time data into 1 ms bins, and calculated the mean dwell time two different ways: (1) using the maximum likelihood estimator (MLE, unbinned) (2) fitting the binned data, but excluding the lowest-duration bins. Figure s1 and table s1 show the results for this analysis. Both the MLE and fitting to the histogram with no bins removed estimate the mean dwell time to be $\approx$ 3.5 $ms$, whereas removing the first bin from the fitting results in a mean dwell time of $\approx$ 2 $ms$ and the result is stable regardless of the number of bins removed. This suggests that fits can be appropriately performed by simply excluding the bin from 0-1 ms. We estimate the loss fraction from the data as

$$loss\ fraction\ (0-1\ ms) = 1 - \frac{N_{obs}(0-1ms)}{N_{expected}(0-1ms)} \tag{s2}$$

where $N_{expected}$ is model-dependent and calculated from the fits excluding the 0-1 ms data. We estimate that the loss fraction between 0-1 ms is 0.80 $\pm$ 0.17. This value is systematically over-estimated by virtue of assuming the single-exponential distribution function. Any deviation from this model, such as further hidden reverse processes in the f|b step, will necessarily lead to longer dwell times and therefore to an overestimate of the loss fraction.

We also considered the weighted MLE:

$$\langle t \rangle = \frac{\Sigma_i w_i t_i}{\Sigma_i w_i} \tag{s3}$$

with weights given by

$$w_i = \begin{cases} (1 - loss\ fraction)^{-1} & t < 1\ ms \\ 1 & t > 1\ ms \end{cases} \tag{s4}$$

Table s1 shows the weighted MLE as a function of the loss fraction. For a loss fraction of 0.6, our weighted MLE estimate of the mean is nearly identical to the unweighted MLE, which we know must be an overestimate of the mean dwell time. For a loss fraction of 0.99 the estimated dwell time is substantially underestimated. We choose a loss fraction of 0.93, which gives a reasonable estimate of the size of the effect and calculated means using the weighted MLE. Ultimately, above a mean dwell time of $\approx$ 5 $ms$ the correction is negligible.

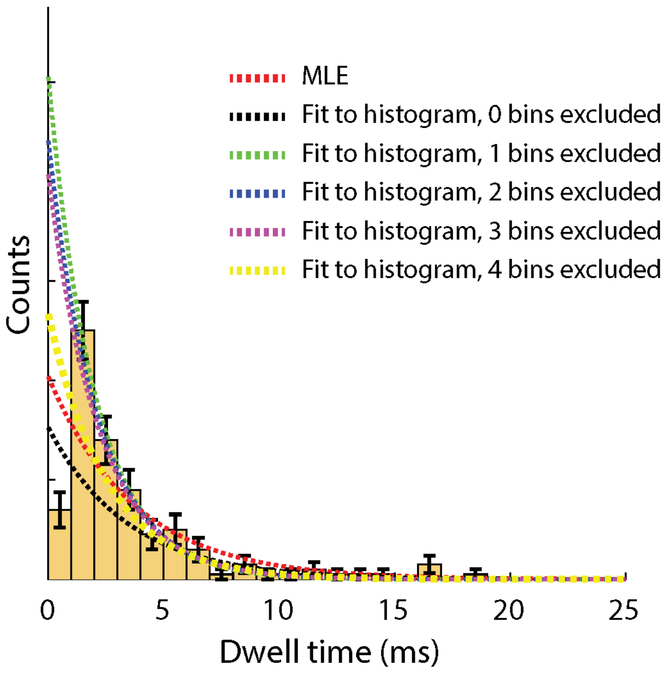


FIGURE S1. Distribution of f|b dwell times for the [ATP]-dependent state in the 5′ force-opposing configuration. Dashed lines indicated different methods of fitting the data: (red) Maximum-likelihood estimator (black) fit to histogram with no bins removed (green) fit to histogram with data < 1 ms excluded (blue) fit to histogram with data < 2 ms excluded (pink) fit to histogram with data < 3 ms excluded (yellow) fit to histogram with data < 4 ms excluded.

| Method | $\langle t \rangle_{est}(ms)$ |
|---|---|
| MLE | $3.7 \pm 0.3$ |
| Histogram all bins | $3.5 \pm 0.3$ |
| Histogram exclude 0-1 ms | $2.0 \pm 0.1$ |
| Histogram exclude 0-2 ms | $2.1 \pm 0.1$ |
| Histogram exclude 0-3 ms | $2.2 \pm 0.2$ |
| Histogram exclude 0-4 ms | $2.5 \pm 0.3$ |
| Weighted MLE (loss fraction 0-1 ms = 0.6) | $3.4 \pm 0.3$ |
| Weighted MLE (loss fraction 0-1 ms = 0.7) | $3.2 \pm 0.3$ |
| Weighted MLE (loss fraction 0-1 ms = 0.8) | $2.9 \pm 0.3$ |
| Weighted MLE (loss fraction 0-1 ms = 0.9) | $2.4 \pm 0.3$ |
| Weighted MLE (loss fraction 0-1 ms = 0.95) | $1.9 \pm 0.2$ |
| Weighted MLE (loss fraction 0-1 ms = 0.99) | $1.2 \pm 0.1$ |

TABLE S1. Estimated dwell time for f|b [ATP]-dependent states with single exponential fits based on different methods.

1.9. **DNA design.** Figure s2 shows the DNA sequences used in this study. RecQ preferentially loads at a ssDNA/dsDNA junction on the $3'$ end, and translocates from $3'$ to $5'$.

In the first design we use a hairpin sequence with a 30-nt DNA duplex with a Y-tail, at which RecQ loads. The $5'$ end has a cholesterol tag to facilitate insertion of the $5'$ end into the bilayer. The $3'$ end is then drawn into the pore, which is encouraged by the presence of a phosphate on the $3'$ end of the DNA.

The second design has the same sequence at the first, but instead of a hairpin sequence we use a short adapter that is annealed to the template. This adapter has a $5'$ cholesterol to facilitate insertion into the bilayer. Because the sequence being measured in MspA is displaced from the enzyme by $\approx 15 - 20\ nt$, when the 'measure-sequence' is in MspA, the entirety of the short duplex must have dissociated, leaving us with force-assisted ssDNA translocation. A second version of this design has a longer measure-sequence (19-nt) and has a further 33-nt poly-T sequence on the $5'$ end of the template. This controls for the potential effects of sequence-dependent kinetics.

The final design is for force-opposed translocation. We again anneal a short blocker to encourage RecQ binding to the DNA, but in this configuration we have a $3'$ cholesterol on both the template and blocker sequences. We include a 25-nt poly-T sequence adjacent to the measure-sequence, ensuring that poly-T is in the RecQ when the measure sequence is in the pore. This ensures that the same sequence is in RecQ for both force-assisting and force-opposing ssDNA translocation.

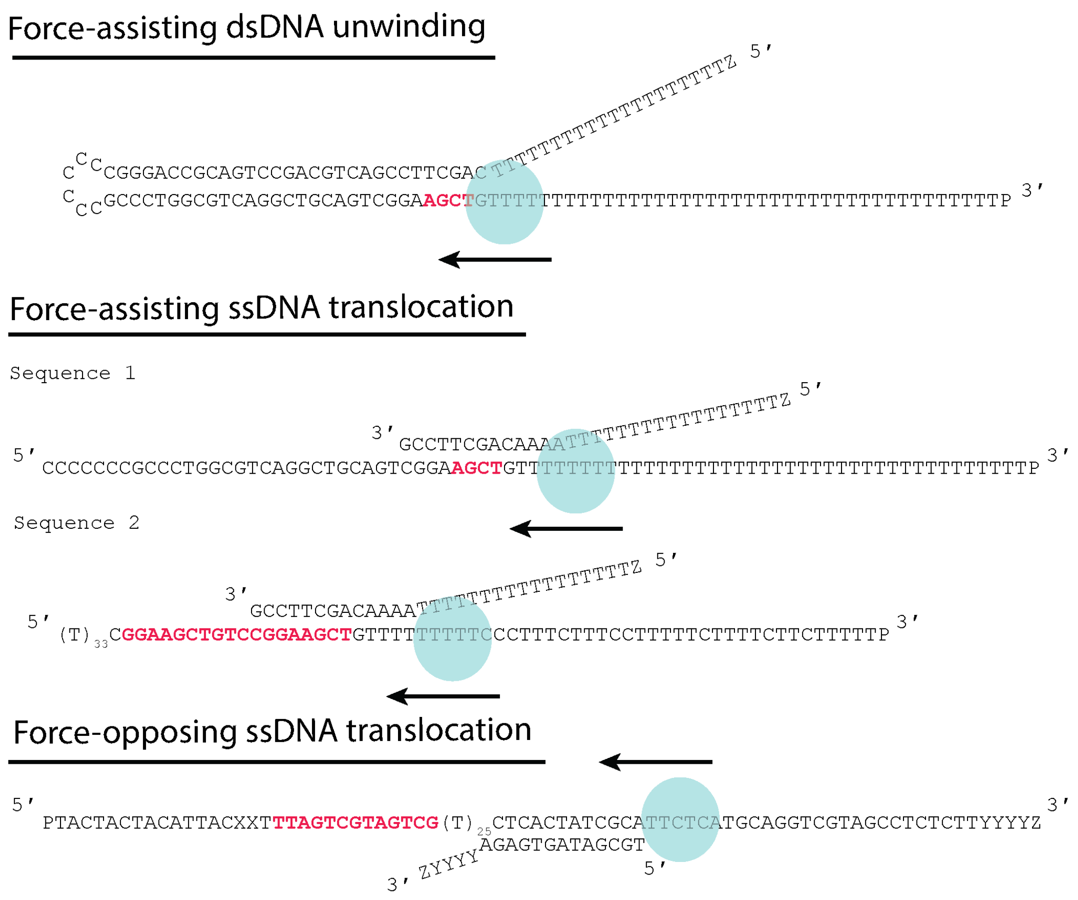


FIGURE S2. DNA designs used in this study. Letters in red are the ‘measure-sequence’, the nucleotides in the MspA constriction when kinetic measurements are made. P is a phosphate to encourage entry into the nanopore, X is an abasic, Y is an 18-carbon spacer and Z is a cholesterol tag to bind DNA to the lipid bilayer. The blue circle represents the RecQ binding site and the arrow the direction of helicase motion on the template. (Top) Force-assisted unwinding of a hairpin. (Middle) Force-assisting ssDNA translocation (Bottom) Force-opposing ssDNA translocation.

##### 1.10. **Experimental Statistics.**

| DNA configuration | $[ATP](\mu M)$ | $[ADP](\mu M)$ | $[ATP\gamma S](\mu M)$ | $Voltage(mV)$ | $N_{event}$ |
|---|---|---|---|---|---|
| dsDNA $f_{assist}$ | 10 | 0 | 0 | 180 | 27 |
| dsDNA $f_{assist}$ | 20 | 0 | 0 | 180 | 106 |
| dsDNA $f_{assist}$ | 50 | 0 | 0 | 180 | 30 |
| dsDNA $f_{assist}$ | 250 | 0 | 0 | 180 | 21 |
| dsDNA $f_{assist}$ | 1000 | 0 | 0 | 180 | 144 |
| dsDNA $f_{assist}$ | 2000 | 0 | 0 | 180 | 31 |
| dsDNA $f_{assist}$ | 10 | 10 | 0 | 180 | 29 |
| dsDNA $f_{assist}$ | 20 | 20 | 0 | 180 | 27 |
| dsDNA $f_{assist}$ | 50 | 50 | 0 | 180 | 35 |
| dsDNA $f_{assist}$ | 250 | 250 | 0 | 180 | 21 |
| dsDNA $f_{assist}$ | 1000 | 1000 | 0 | 180 | 26 |
| dsDNA $f_{assist}$ | 2000 | 2000 | 0 | 180 | 29 |
| dsDNA $f_{assist}$ | 10 | 40 | 0 | 180 | 15 |
| dsDNA $f_{assist}$ | 20 | 80 | 0 | 180 | 25 |
| dsDNA $f_{assist}$ | 50 | 200 | 0 | 180 | 31 |
| dsDNA $f_{assist}$ | 250 | 1000 | 0 | 180 | 22 |
| dsDNA $f_{assist}$ | 750 | 2800 | 0 | 180 | 33 |
| dsDNA $f_{assist}$ | 1000 | 0 | 500 | 180 | 53 |
| ssDNA $f_{assist,seq1}$ | 1000 | 0 | 0 | 180 | 41 |
| ssDNA $f_{assist,seq2}$ | 2500 | 0 | 0 | 120 | 55 |
| ssDNA $f_{assist,seq2}$ | 2500 | 0 | 0 | 140 | 51 |
| ssDNA $f_{assist,seq2}$ | 2500 | 0 | 0 | 160 | 43 |
| ssDNA $f_{assist,seq2}$ | 2500 | 0 | 0 | 180 | 84 |
| ssDNA $f_{assist,seq2}$ | 2500 | 0 | 0 | 200 | 27 |
| ssDNA $f_{assist,seq2}$ | 2500 | 0 | 0 | 220 | 24 |
| ssDNA $f_{oppose}$ | 2500 | 0 | 0 | 120 | 60 |
| ssDNA $f_{oppose}$ | 2500 | 0 | 0 | 140 | 47 |
| ssDNA $f_{oppose}$ | 2500 | 0 | 0 | 160 | 78 |
| ssDNA $f_{oppose}$ | 2500 | 0 | 0 | 180 | 84 |
| ssDNA $f_{oppose}$ | various† | various† | 0 | 180 | 100 |
| ssDNA $f_{oppose}$ | 2500 | 0 | 0 | 200 | 39 |
| ssDNA $f_{oppose}$ | 2500 | 0 | 0 | 220 | 36 |

TABLE s2. Table of experimental statistics. DNA sequences are shown in Figure s2. † The purpose of these experiments was to determine which of the two steps responded to changes in ADP, and contains data from several different experiments: $[ATP]/[ADP] = 2500/1500$, 1500/2500 and 1000/3000. These data were not used in any capacity that required exact values of the concentrations. They were solely used for identification of [ATP]-dependent states in force opposing configuration (Fig s5).

#### 2. Competitive inhibition model for [ATP]-dependent state

From our kinetic model for the [ATP]-dependent state (Fig. 3A, MT), it can be shown that the mean $f|f$ dwell time is given by (Craig et al. 2017):

$$\langle t\rangle_{f|f} = \frac{K + d\cdot[ADP] + [ATP]}{V\cdot[ATP]}. \tag{s5}$$

where $V$ is the velocity of the [ATP]-dependent state at saturating [ATP], $K$ is the [ATP] at which the velocity is half of its maximum, and $d = K/K_I$ with $K_I$ being the inhibition equilibrium constant. Fitting this expression to the data in Figure s3 yields $V = 106 \pm 7\ s^{-1}$, $K = 39 \pm 5\ \mu M$ and $d = 0.77 \pm 0.13$. From these and the measured value of $k_D = 220 \pm 10\ s^{-1}$ we calculate $k_{-D} = \frac{d\cdot k_D}{K} = 4.3 \pm 0.9\ s^{-1}\cdot\mu M^{-1}$.

The simplest possible alternative model with a single ATP and ADP binding step is shown in figure s3. This model is similar to those found in DNA polymerases (Lieberman *et al.* 2013 JACS) in which the enzyme oscillates between pre- and post-translocation states before NTP binding rectifies forwards motion. The average dwell time as a function of [ATP] and [ADP] in this mechanism is:

$$\langle t\rangle_{f|f} = \frac{K + d\cdot[ADP] + [ATP] + e\cdot[ATP]\cdot[ADP]}{V\cdot[ATP]}. \tag{s6}$$

In this alternative model, the parameter $e$ couples the averages dwell time to the product $[ATP]$ and $[ADP]$. In this mechanism, simultaneously increasing ATP and ADP results in shifting the chemical equilibrium away from the conformational change steps ($k_{\pm1}$ and $k_{\pm2}$), thereby *disfavoring* conformational changes and prolonging the f|f dwell time. In contrast, for our competitive inhibition mechanism (Fig 3A.) simultaneously increasing ATP and ADP shifts the chemical equilibrium in a way that *favors* conformational changes. Fitting s6 to the data yields the same estimates for $V$, $K$ and $d$ as above with $e = (0.5 \pm 1.5)\cdot 10^{-4}\cdot\mu M^{-1}$. This value of $e$ is consistent with 0, which is an unphysical result, since it requires that both $k_H$ and $k_{-H}$ be simultaneously 0 (Craig *et al.* 2017 PNAS).

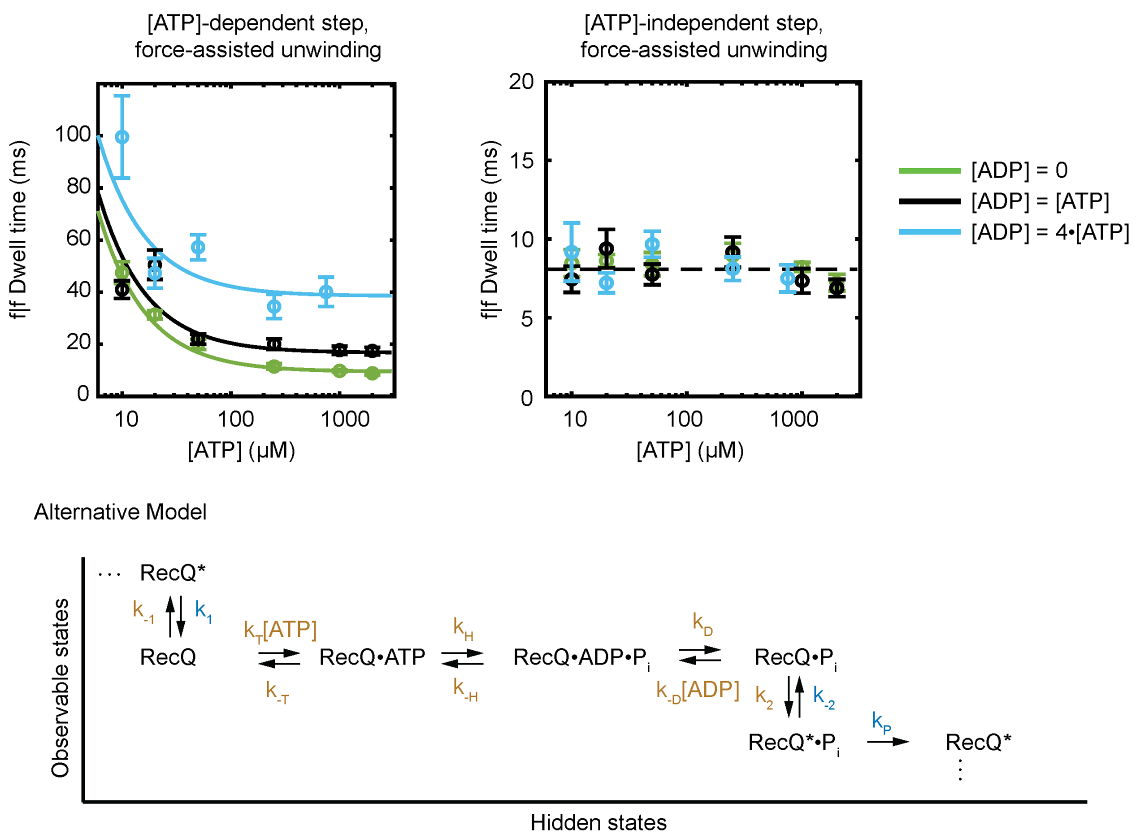


FIGURE S3. (Top, Left) $f|f$ dwell time vs. [ATP] for the [ATP]-dependent state with the ratio of [ADP]:[ATP] held fixed at 0 (green) 1:1 (black) and 1:4 (light blue). Solid lines are simultaneously fit based on equation s5. (Top, Right) $f|f$ dwell time vs. [ATP] for the [ATP]-independent state. Colors are the same as left. The black dashed line is the mean taken over all data. (Bottom) An alternative model of ATPase activity.

#### 3. Analysis of sequence-dependent effects on ATP$\gamma$S hydrolysis

In M.T. Figure 2 we observed that ATP$\gamma$S inhibition of RecQ had two distinct behaviors depending on the sequence position: a hydrolysis mechanism in which RecQ hydrolyzes ATP$\gamma$S and an ejection mechanism in which RecQ oscillates between open and closed conformations, ejecting ATP$\gamma$S during the open state. Upon ATP binding RecQ then proceeded through its standard ATP hydrolysis pathway. We sought to determine which rate constants in our kinetic models were being effected by the DNA sequence.

Figure s4 shows our mechanism for the [ATP]-independent step during ATP$\gamma$S hydrolysis. Under the assumption that ATP$\gamma$S hydrolysis is the rate-limiting we can write the following expressions:

$$p_{b|f,\gamma} = \frac{k_{-S\gamma}}{k_{-S\gamma} + k_{H\gamma}} \tag{s7}$$

$$\tau_{b|f} = \frac{1}{k_{-S\gamma} + k_{H\gamma}} \tag{s8}$$

where $p_{b|f,\gamma}$ is the probability of a backwards step given an ATP$\gamma$S binding event, $\tau_{b|f}$ is the conditional dwell time of the backwards steps, $k_{H\gamma}$ is the rate of ATP$\gamma$S hydrolysis and $k_{-S\gamma}$ is the rate of RecQ opening when ATP$\gamma$S is bound. We can then rearrange the above expressions to solve for $k_{-S\gamma}$ and $k_{H\gamma}$:

$$k_{-S\gamma} = \frac{p_{b|f,\gamma}}{\tau_{b|f}} \tag{s9}$$

$$k_{H\gamma} = \frac{1 - p_{b|f,\gamma}}{\tau_{b|f}}. \tag{s10}$$

By measuring $p_{b|f,\gamma}$ and $\tau_{b|f}$ we can calculate the rate constants at each sequence position. For positions 2 and 3 we have enough measurements of the dwell time to measure $\tau_{b|f}$ directly. For positions 1 and 4 we do not, but if we know that ATP$\gamma$S hydrolysis is the rate-limiting behavior we can use the fact that $f|f$ dwell times longer than 50 $ms$ are almost certainly ATP$\gamma$S hydrolysis events to estimate

$$\tau_{b|f} = \tau_{f|f,ATP\gamma S} \approx \tau_{f|f,>50\ ms}. \tag{s11}$$

For DNA position 1 we have $\tau_{f|f,>\ 50\ ms} = 1.0 \pm 0.2\ s$ and $p_{b|f} < 0.05$ (0 backwards steps in 22 ATP$\gamma$S binding events). We therefore estimate $k_{-S\gamma} < 0.05\ s^{-1}$ and $k_{H\gamma} = 1.0 \pm 0.2\ s^{-1}$. For DNA position 2 we have $\tau_{b|f} = 0.17 \pm 0.06\ s$ and $p_{b|f,\gamma} = 0.94 \pm 0.04$ and therefore estimate $k_{-S\gamma} \approx 5 \pm 1\ s^{-1}$ and $k_{H\gamma} = 0.35 \pm 0.1\ s^{-1}$. These results suggest that the sequence-dependent effects manifest primarily by enhancing the rate of backwards steps, with a factor of at least 100 difference between estimated values of $k_{-S\gamma}$ between positions 1 and 2, whereas estimated values of $k_{H\gamma}$ are much more similar.

These experiments were done at $V = 180\ mV$ assisting force. We use the measured value of the force-coupling parameter $\alpha_{-SD} \approx 0.014\ mV^{-1}$ to extrapolate the value of $k_{-S\gamma}$ to zero force (see section s7):

$$k_{-S\gamma}(V = 0) = k_{-S\gamma}(V = 180\ mV) \cdot exp(\alpha_{-SD} \cdot V) \approx 10 \cdot k_{-S\gamma}(V = 180\ mV). \tag{s11}$$

Applying this expression to positions 1 and 2 from above we calculate at position 1 that $k_{-S\gamma}(V = 0) < 0.5\ s^{-1}$ and at position 2 that $k_{-S\gamma}(V = 0) \approx 50\ s^{-1}$. Because $k_{ST}(V = 0) \approx 350\ s^{-1}$ we have $k_{-S\gamma}/k_{S\gamma} < 1/7$ at zero force ($k_{ST}$ is the closing rate when ATP is bound, table s5). Therefore in the absence of force RecQ primarily is in the [ATP]-independent state while bound to ATP$\gamma$S, corresponding to closed RecA-like domains.

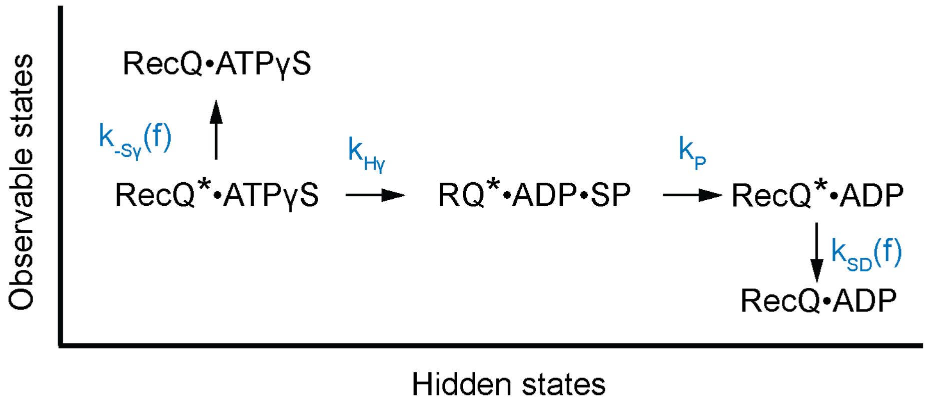


Figure s4. Kinetic Mechanism used for analysis of ATP$\gamma$S hydrolysis. $k_{H\gamma}$ is the rate of ATP$\gamma$S hydrolysis.

#### 4. Identification of [ATP]-dependent states in force-opposing configuration

We identified [ATP]-dependent states in the force-opposing configuration by performing experiments at saturating [ATP] in the presence and absence of ADP. Figure s5 shows the dwell time and probability of a backwards step in these conditions. The average dwell time increases slightly at half-integer positions, and the probability of a backwards step increases substantially at these same positions 3. We identify the half-integer positions as [ADP]-dependent, and therefore as [ATP]-dependent.

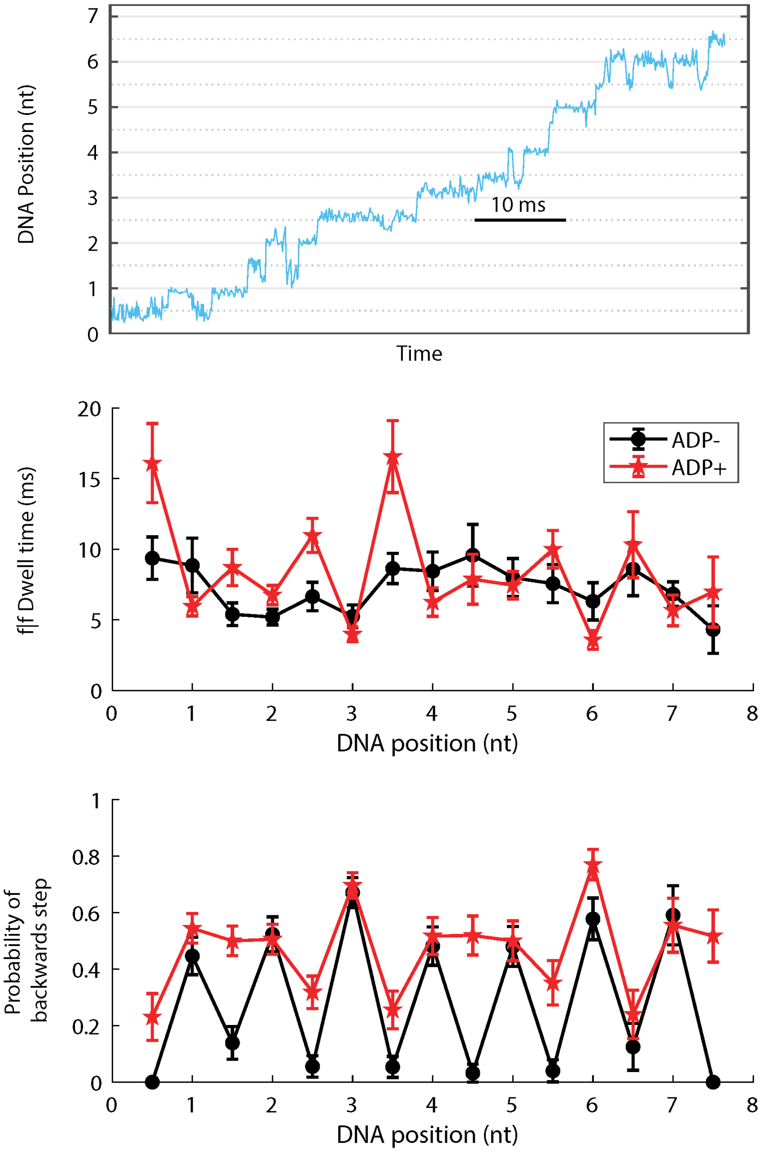


Figure s5. (Top) Position vs. time trace for RecQ translocation on ssDNA against 180 mV opposing force at saturating $[ATP] = 2500\ \mu M$. (Middle) $f|f$ dwell time vs. DNA position for RecQ translocation on ssDNA at 180 mV opposing force in the absence of ADP (black) and the presence of equimolar ADP (red). (Bottom) Probability of a backwards step vs. DNA position for the same conditions as above.

| Parameter | [ATP]-independent ($[ADP] = 0$) | [ATP]-independent ($[ADP] \approx [ATP]$) | [ATP]-dependent ($[ADP] = 0$) | [ATP]-dependent ($[ADP] \approx [ATP]$) |
|---|---|---|---|---|
| $\langle t \rangle_{f\|f} (ms)$ $f = +37\ pN$ | $7.3 \pm 0.3$ | $7.1 \pm 0.4$ | $8.5 \pm 0.4$ | $17 \pm 1$ |
| $p_{b\|f}$ $f = +37\ pN$ | $0.003 \pm 0.002$ | 0 | $0.008 \pm 0.004$ | $0.005 \pm 0.005$ |
| $\langle t \rangle_{f\|f} (ms)$ $f = -37\ pN$ | $7.0 \pm 0.5$ | $6.1 \pm 0.3$ | $7.4 \pm 0.5$ | $10.1 \pm 0.6$ |
| $p_{b\|f}$ $f = -37\ pN$ | $0.51 \pm 0.02$ | $0.53 \pm 0.02$ | $0.07 \pm 0.02$ | $0.37 \pm 0.03$ |

TABLE S3. Kinetic measurements comparing the effects of ADP on kinetic parameters of the [ATP]-dependent and [ATP]-independent states in assisting and opposing force experiments.

#### 5. Mechanism of $b|f$ [ATP]-independent steps; ruling out a futile hydrolysis pathway

We previously observed in Superfamily 2 helicase Hel308 that $b|f$ [ATP]-independent states were consistent with a two-pathway mechanism: one of which was an ‘immediate reversal’ pathway in which the conformational closure of the helicase RecA-like domains is followed by an opening, and a ‘futile hydrolysis’ mechanism in which ATP was hydrolyzed, but instead of stepping forwards, the helicase stepped backwards to the preceding [ATP]-dependent state (Fig. s6A, Craig *et al.* 2017 PNAS, Craig *et al.* 2021 Essays in Biochemistry). In Hel308 most backwards steps followed the futile hydrolysis pathway. Here we analyze conditional dwell time distributions to determine whether futile hydrolysis occurs during RecQ translocation on DNA.

Figure s6B shows the dwell time distributions for $f|f$ and $f|b$ [ATP]-dependent states. The $f|f$ steps (the ‘standard step’) are consistent with multiple rate-limiting processes, whereas the $f|b$ steps are substantially faster than the $f|f$ steps and more consistent with a single-exponential distribution, implying that the $f|b$ steps are a single step corresponding to the closing of ATP-bound RecQ.

If we assume that the futile hydrolysis pathway is the dominant back-stepping pathway, then the initial conditions for $f|f$ and $f|b$ [ATP]-dependent states would be identical, because both initial conditions are an ADP-bound open conformation of the helicase. However, if this were the case, we would then expect the distribution functions to be identical. Instead we observe a multi-pathway process for the $f|f$ steps and a single process for $f|b$ steps, suggesting the ‘immediate reversal’ pathway is dominant.

From this result, we then compared the distribution of $f|f$ [ATP]-independent steps to the distribution of $b|f$ [ATP]-independent steps. The distributions are very similar, with a slight bias towards short dwell times in the $b|f$ steps. This implies that the rate-limiting step of the [ATP]-independent step is ATP hydrolysis, consistent with quantitative analysis our kinetic model (Fig. s12, section s8) which found $k_H \approx 125\ s^{-1}$ and $k_{SD}(V = -180\ mV) \approx 400\ s^{-1}$.

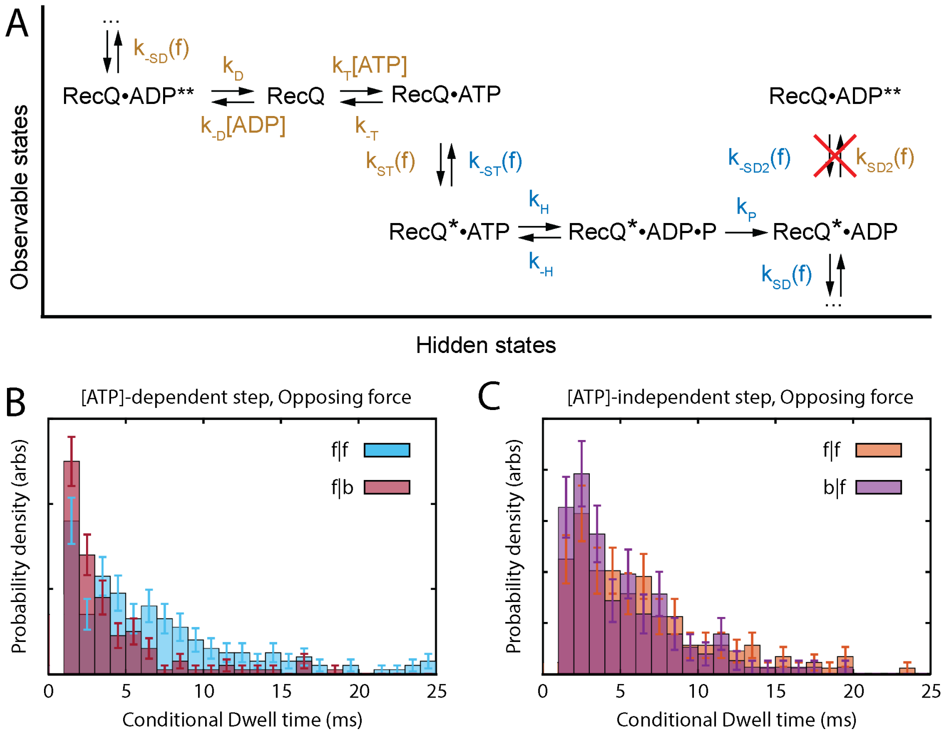


Figure s6. (A) Possible futile hydrolysis mechanism for RecQ. (B) Conditional dwell time distribution for $f|f$ (blue) and $f|b$ (burgandy) [ATP]-dependent states at 180 mV opposing force. (C) Conditional dwell time distribution for $f|f$ (orange) and $b|f$ (purple) [ATP]-independent steps at 180 mV opposing force.

#### 6. Step size measurements in force-assisting and force-opposing ssDNA translocation experiments

We measure the step size as in (Derrington *et al.* 2015, Nature Biotechnology). Briefly, we identify [ATP]-dependent and [ATP]-independent steps, and calculate the spline curve for both steps independently. We then calculate the distance by finding the phase shift, $\varphi$ (measured in nt), that minimizes the sum square difference between the two curves. To calculate an error bar, we vary the values of the ion-current used to construct the spline by adding gaussian noise with width given by the s.e.m. and repeat the calculation 100 times, taking the standard deviation at the end (Figure s7). We find $\varphi_{3'} = 0.56 \pm 0.04\ nt$ and $\varphi_{5'} = 0.62 \pm 0.02\ nt$.

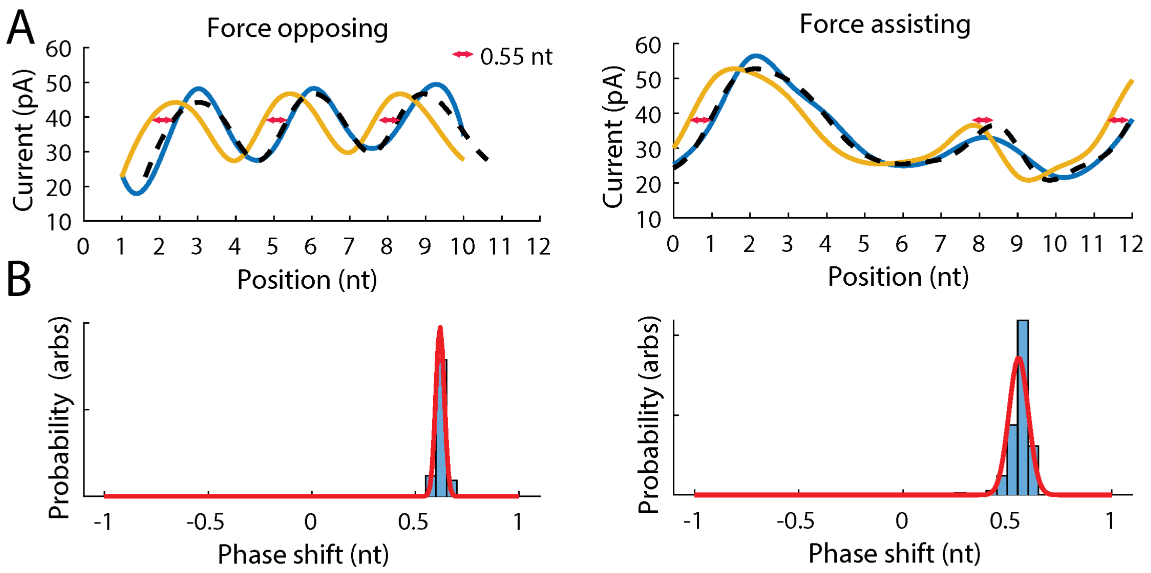


Figure s7. (A) Step size measurements in force opposing (left) and force-assisting (right) configurations. The [ATP]-dependent state is in gold and the [ATP]-independent state in blue. The black dashed line shows the [ATP]-dependent state offset by the best-fit value of the step size. (B) Monte-carlo distribution of fit values for the step-size.

#### 7. Modelling force-dependent rate constants in SPRNT; a constraint equation

We apply single-molecule kinetics theory together with transition state theory to interpret the results of force-dependent translocation on ssDNA. Figure s8 shows a two-state energy diagram, with two energy minima separated by a distance $x_s$ and a transition state, which is separated from the left-most minimum by a distance $x_t$. To first-order (Walcott 2008 J. Chem. Physics) the force dependence of the forwards and reverse rate constants is:

(s12a) $$k_+(f) = k_{+,0} \cdot exp(f \cdot x_t/kT)$$

(s12b) $$k_-(f) = k_{-,0} \cdot exp(f \cdot (x_s - x_t)/kT).$$

In SPRNT we apply force via the voltage. We define $\lambda$ to be the linear charge density of ssDNA so that the force on a DNA molecule being moved through the voltage V is

(s13) $$f = \lambda \cdot V.$$

The maximal charge density of ssDNA is $\lambda = 1e^- \cdot nt^{-1}$. We then define a new parameter $\alpha_1$ as

(s14) $$\alpha_1 \equiv \frac{\lambda}{kT} = \frac{1e^- \cdot nt^{-1}}{26\ meV} = 0.039\ mV^{-1} \cdot nt^{-1}$$

In SPRNT, we measure the coupling between rate and voltage in the context of a single physical step as

(s15) $$k_i(V) = k_{V=0} \cdot exp(\alpha_i \cdot V)$$

where $\alpha_i = \alpha_1 \cdot x_{t,i}$ is a measurable parameter in SPRNT, representing the coupling between a kinetic rate corresponding to a physical step and the voltage. Because $\alpha_1$ assumes a maximal charge density for ssDNA, it also sets a maximum on the extent to which voltage can couple to enzyme kinetics. For a 1-nt step composed of n sub-nt steps, with distances to the transition state $x_{t,i}$ we necessarily have

(s16) $$\Sigma_{i=1}^{2 \cdot n} |x_{t,i}| = 1 \cdot nt$$

Multiplying this by $\alpha_1$ we find

(s17) $$\Sigma_{i=1}^{2 \cdot n} |\alpha_i| < \alpha_1.$$

This is a physical upper limit on the effect that voltage can have on force-dependent kinetics in Nanopore Tweezers. Because RecQ moves DNA through MspA in two sub-nt steps we have n = 2 and four force-dependent rate constants.

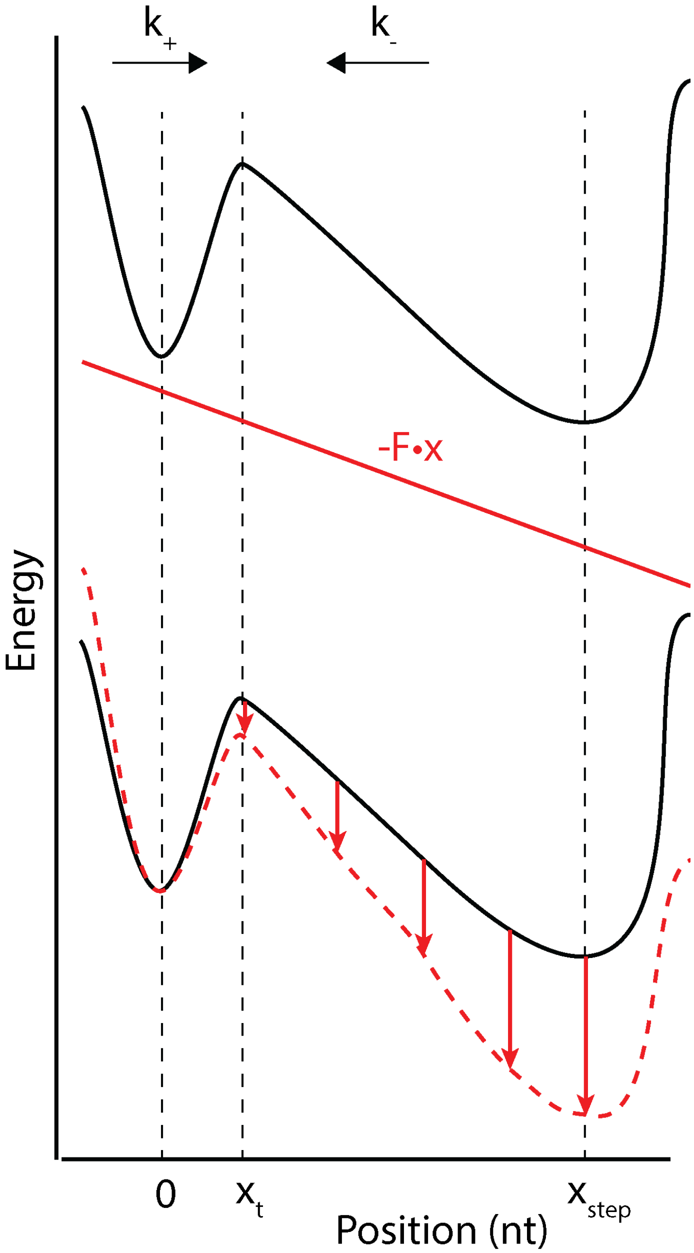


FIGURE S8. Example energetic landscape for a physical step by an enzyme. The first energy minimum is given a physical coordinate $x = 0\ nt$ by convention. The transition state is located at coordinate $x = x_t$ and the post-translocated state is at $x = x_{step}$, which can be measured in SPRNT. We then infer the position of the transition state $x_t$ based on how the enzyme responds to a mechanical force.

#### 8. Parameter estimation in force-dependent RecQ translocation on ssDNA

In our kinetic model of RecQ translocation (Fig. s9) we have four force-dependent rate constants: $k_{SD}$, $k_{-SD}$, $k_{ST}$ and $k_{-ST}$. By convention, we define the $\alpha_i$ for these parameters to be strictly greater than 0 and include negative signs in the exponential terms for backwards transitions. Because the distance from the [ATP]-dependent state to the [ATP]-independent state is similar for the force-assisting and force-opposing geometries (Fig. s7), we assume that the $\alpha_i$ do not change between the orientations. The expressions for these four rate constants are:

(s18a) $$k_{SD}(V) = k_{SD} \cdot e^{\alpha_{SD} \cdot V}$$

(s18b) $$k_{-SD}(V) = k_{-SD} \cdot e^{-\alpha_{-SD} \cdot V}$$

(s18c) $$k_{ST}(V) = k_{ST} \cdot e^{\alpha_{ST} \cdot V}$$

(s18d) $$k_{-ST}(V) = k_{-ST} \cdot e^{-\alpha_{-ST} \cdot V}.$$

We used conditional dwell times for both the [ATP]-dependent and [ATP]-independent states, together with the average velocity to estimate kinetics parameters. Our force-dependent experiments were done in the absence of ADP and phosphate, and at saturating [ATP]. We therefore take the limits $[ADP] \to 0$, $[P_i] \to 0$ and $[ATP] \to \infty$. In addition, we assume that $k_{-H} \ll k_p$, effectively making ATP hydrolysis irreversible. Using the master equation we write the connection matrices for the [ATP]-dependent state and [ATP]-independent state as:

(s19) $$M_{[ATP]-dep} = \begin{bmatrix} 0 & k_{-SD}(V) & 0 & 0 & 0 \\ 0 & -k_{-SD}(V) - k_D & k_{-D} \cdot [ADP] & 0 & 0 \\ 0 & k_D & -k_{-D} \cdot [ADP] - k_T \cdot [ATP] & k_{-T} & 0 \\ 0 & 0 & k_T \cdot [ATP] & -k_{-T} - k_{ST}(V) & 0 \\ 0 & 0 & 0 & k_{ST}(V) & 0 \end{bmatrix}$$

(s20) $$M_{[ATP]-indep} = \begin{bmatrix} 0 & k_{-ST}(V) & 0 & 0 & 0 \\ 0 & -k_{-ST}(V) - k_H & 0 & 0 & 0 \\ 0 & k_H & -k_p & 0 & 0 \\ 0 & 0 & k_p & -k_{SD}(V) & 0 \\ 0 & 0 & 0 & k_{SD}(V) & 0 \end{bmatrix}.$$

The initial conditions for the master equation are

(s21a) $$\vec{p}(t=0)_{f|f} = (0,1,0,0,0)^T$$

(s21b) $$\vec{p}(t=0)_{b|f} = (0,1,0,0,0)^T$$

(s21c) $$\vec{p}(t=0)_{f|b} = (0,0,0,1,0)^T.$$

The master equation is then solved for the mean conditional dwell times and probability of a backwards step (Craig *et al.* 2021 Essays in Biochemistry). For the [ATP]-dependent state in the limits described above the expressions are:

(s22a) $$p_{b|f} = \frac{k_{-SD}(V)}{k_{-SD}(V) + k_D}$$

(s22b) $$\langle t \rangle_{f|f} = \frac{k_D + k_{ST}(V) + k_{-SD}(V)}{k_{ST}(V)(k_D + k_{-SD}(V))}$$

(s22c) $$\langle t \rangle_{b|f} = \frac{1}{k_{-SD}(V) + k_D}$$

(s22d) $$\langle t \rangle_{f|b} = \frac{1}{k_{ST}(V)}$$

and for the [ATP]-independent state:

(s23a) $$p_{b|f} = \frac{k_{-ST}(V)}{k_{-ST}(V) + k_H}$$

(s23b) $$\langle t \rangle_{f|f} = \frac{1}{k_{-ST}(V) + k_H} + \frac{1}{k_p} + \frac{1}{k_{SD}(V)}$$

(s23c) $$\langle t \rangle_{b|f} = \frac{1}{k_{-ST}(V) + k_H}$$

(s23d) $$\langle t \rangle_{f|b} = \frac{1}{k_{SD}(V)}.$$

The average velocity is:

(s24) $$velocity = \frac{1}{\langle t \rangle} = [\frac{k_{ST}(V) + k_H + k_{-ST}(V)}{k_{ST}(V) \cdot k_H} + \frac{k_{SD}(V) + k_D + k_{-SD}(V)}{k_{SD}(V) \cdot k_D} + \frac{1}{k_p}]^{-1}.$$

We have measurements and errors for the left hand side of s22-s24. Our goal is to evaluate the parameters on the R.H.S. of these expressions. Our model has 11 free parameters, 24 measurements of the probability of a backwards step ([ATP]-dependent, [ATP]-independent states, at 6 assisting forces and 6 opposing forces), 24 measurements of the $f|f$ dwell time, 12 measurements of the $f|b$ dwell times (opposing force only, due to low probability of backwards step in assisting force configuration), 12 measurements of the $b|f$ dwell time (opposing force only) and 12 velocity measurements. We construct the log-likelihood function as a sum of chi-square functions:

(s25) $$\mathcal{L}(\vec{\theta}|data) = -\Sigma_{i=1}^{9} \chi_i^2.$$

The sum is taken over all observables s22-s24 (4 observables each for the [ATP]-dependent and [ATP]-independent steps, as well as the total velocity). $\vec{\theta}$ is an 11-component parameter vector, and the $\chi^2$ is determined from measurements $x_j$ and errors $\sigma_j$ as

(s26) $$\chi_i^2(\vec{\theta}) = \Sigma_j (\frac{x_j - f_i(\vec{\theta})}{\sigma_j})^2.$$

We use the matlab function fminsearch to minimize the cost function (in this case, $-\mathcal{L}$) starting from an initial guess $\vec{\theta}_0$. Because high-dimensional parameter estimation can have local maxima, we tried 15 initializations of $\vec{\theta}_0$. The parameters tended to converge to the same value as long as physically realistic values (e.g., rate constants of order 1/(mean duration) $\approx$ 100 $s^{-1}$) of the parameters were chosen (Fig. s10-s11, Table s4), suggesting a well-behaved likelihood function near the global maximum. The solutions that were started at physically unrealistic values in some instances converged to values whose $\mathcal{L}$ was much smaller than those started at more realistic values, reflecting poor solutions. To estimate parameters and errors, we chose the $\vec{\theta}_0$ which led to the highest likelihood solution (Trial 15), and performed 100 Monte-carlo simulations in which Gaussian noise was added to the measured data points with width given by the standard error in the mean. Our constraint on the $\alpha_i$ is obeyed since $\alpha_{SD} + \alpha_{-SD} + \alpha_{ST} + \alpha_{-ST} = (0.003 + 0.014 + 10^{-5} + 0.014)\ mV^{-1} \approx 0.031\ mV^{-1} < 0.039\ mV^{-1}$.

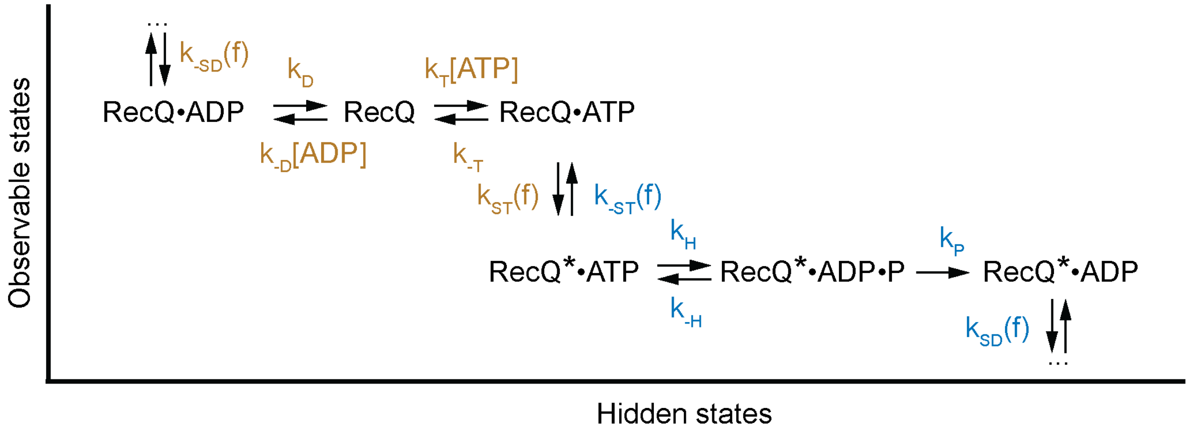


FIGURE S9. Kinetic model of RecQ translocation. ATP and ADP compete for the active site in the [ATP]-dependent state (gold rate constants) before a conformational change results in movements of DNA through the pore. ATP is hydrolyzed during the [ATP]-independent state (blue rate constants), before an ADP-bound conformational change occurs, returning us to the [ATP]-dependent state. ADP is then relased, re-setting the cycle with the enzyme having progressed by 1-nt along the DNA.

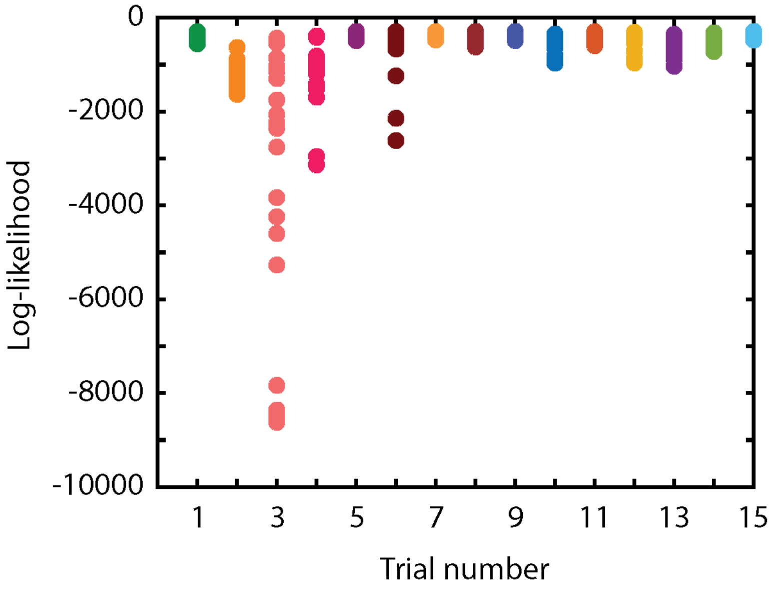


FIGURE S10. Likelihood distribution for parameters $\vec{\theta}$ as a function of the starting co-ordinate $\vec{\theta}_0$. Trials 2-4 were given physically unrealistic starting parameters, leading to low-quality solutions (Table s4).

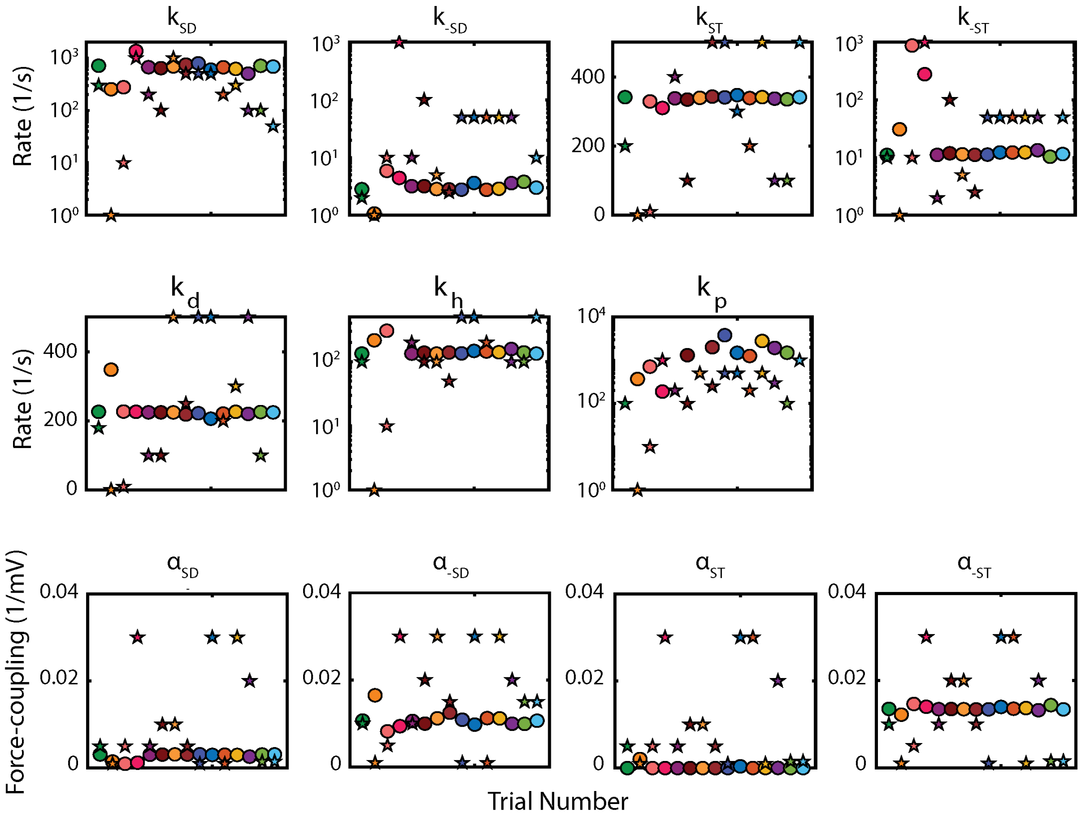


FIGURE S11. Parameter estimation. Each dot represents the median parameter of N = 25 monte carlo trials. Star indicates the start point $\vec{\theta}_0$. Regardless of the start point, parameter estimates are unchanged. Trial numbers correspond to the start values shown in table s4.

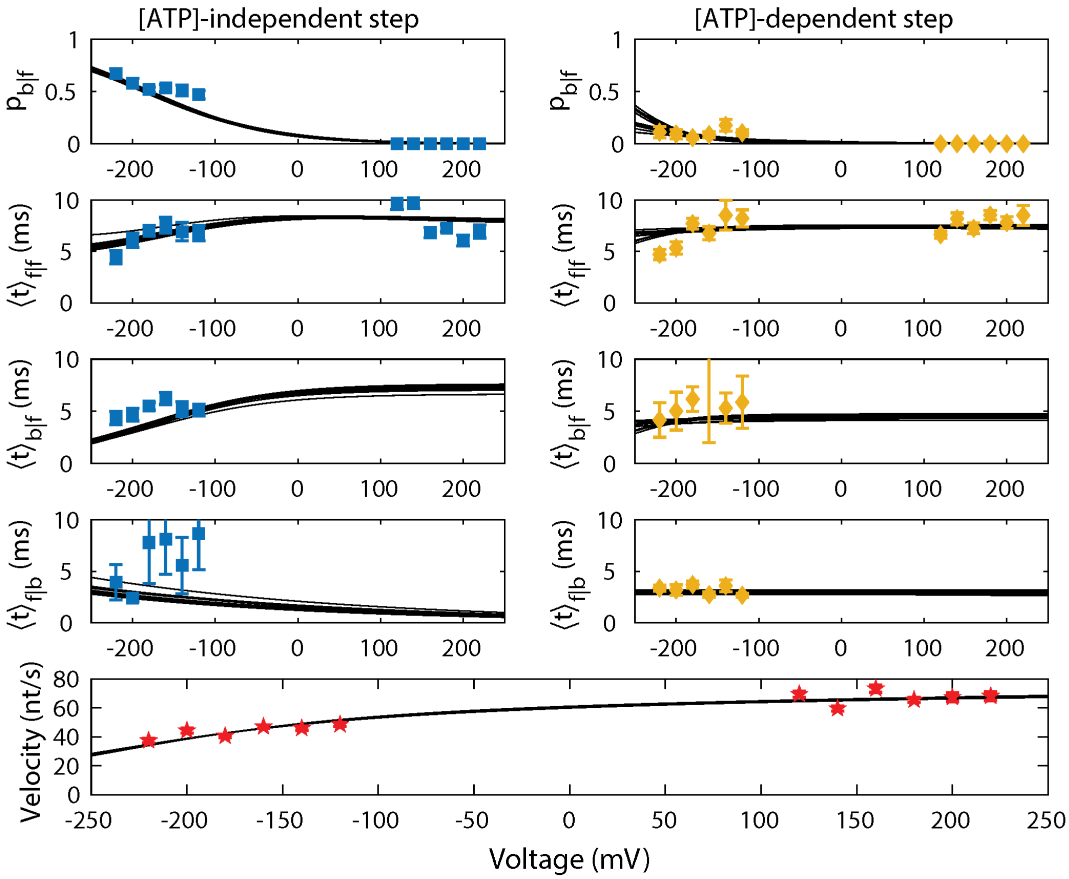


FIGURE S12. Kinetic measurements of RecQ translocation on ssDNA. Negative voltages refer to force-opposing measurements and positive voltages to force-assisting measurements. [ATP]-independent state measurements are in the left column (blue squares) and [ATP]-dependent state measurements are in the right column (gold diamonds). The bottom row is the velocity (red stars). Black lines are fits to the data.

| Trial | $k_{SD}$ | $k_{-SD}$ | $k_{ST}$ | $k_{-ST}$ | $k_D$ | $k_H$ | $k_P$ | $\alpha_{-SD}$ | $\alpha_{-ST}$ | $\alpha_{SD}$ | $\alpha_{ST}$ |
|---|---|---|---|---|---|---|---|---|---|---|---|
| 1 | 300 | 2 | 200 | 10 | 180 | 100 | 100 | 0.01 | 0.01 | 0.005 | 0.005 |
| 2 | 1 | 1 | 1 | 1 | 1 | 1 | 1 | 0.001 | 0.001 | 0.001 | 0.001 |
| 3 | 10 | 10 | 10 | 10 | 10 | 10 | 10 | 0.005 | 0.005 | 0.005 | 0.005 |
| 4 | 1000 | 1000 | 1000 | 1000 | 1000 | 1000 | 1000 | 0.03 | 0.03 | 0.03 | 0.03 |
| 5 | 200 | 10 | 400 | 2 | 100 | 200 | 200 | 0.01 | 0.01 | 0.005 | 0.005 |
| 6 | 100 | 100 | 100 | 100 | 100 | 100 | 100 | 0.02 | 0.02 | 0.01 | 0.01 |
| 7 | 1000 | 5 | 1000 | 5 | 500 | 100 | 500 | 0.03 | 0.02 | 0.01 | 0.01 |
| 8 | 500 | 2.5 | 500 | 2.5 | 250 | 50 | 250 | 0.015 | 0.01 | 0.005 | 0.005 |
| 9 | 500 | 50 | 500 | 50 | 500 | 500 | 500 | 0.001 | 0.001 | 0.001 | 0.001 |
| 10 | 500 | 50 | 300 | 50 | 500 | 500 | 500 | 0.03 | 0.03 | 0.03 | 0.03 |
| 11 | 200 | 50 | 200 | 50 | 200 | 200 | 200 | 0.001 | 0.03 | 0.001 | 0.03 |
| 12 | 300 | 50 | 500 | 50 | 300 | 1000 | 500 | 0.03 | 0.001 | 0.03 | 0.01 |
| 13 | 100 | 50 | 100 | 50 | 500 | 100 | 300 | 0.02 | 0.02 | 0.02 | 0.02 |
| 14 | 100 | 0.1 | 100 | 0.1 | 100 | 100 | 100 | 0.015 | 0.0015 | 0.0015 | 0.0015 |
| 15 | 50 | 10 | 500 | 50 | 1000 | 500 | 1000 | 0.015 | 0.0015 | 0.0015 | 0.0015 |

TABLE S4. Starting values $\vec{\theta}_0$ for optimization of the cost function (s25). All $k_i$ have units of $s^{-1}$ and all $\alpha_i$ have units of $mV^{-1}$.

| Parameter | Unit | Value |
|---|---|---|
| $k_{SD}$ | $s^{-1}$ | $700 \pm 100$ |
| $\alpha_{SD}$ | $mV^{-1}$ | $0.0031 \pm 0.0002$ |
| $k_{-SD}$ | $s^{-1}$ | $2.3 \pm 1.4$ |
| $\alpha_{-SD}$ | $mV^{-1}$ | $0.014 \pm 0.006$ |
| $k_{ST}$ | $s^{-1}$ | $350 \pm 20$ |
| $\alpha_{ST}$ | $mV^{-1}$ | $< 10^{-5}$ |
| $k_{-ST}$ | $s^{-1}$ | $11 \pm 1$ |
| $\alpha_{-ST}$ | $mV^{-1}$ | $0.014 \pm 0.001$ |
| $k_D$ | $s^{-1}$ | $220 \pm 10$ |
| $k_{-D}$ | $s^{-1} \cdot \mu M^{-1}$ | $4.3 \pm 0.9$ |
| $k_H$ | $s^{-1}$ | $136 \pm 7$ |
| $K_T$ | $\mu M$ | $100 \pm 18$ |
| $k_{-T}$ | $s^{-1}$ | $> 1000$ |
| $k_P$ | $s^{-1}$ | $> 1000$ |

TABLE S5. RecQ estimated model parameters as defined Figure 3, the $\alpha_i$ represent the exponential coupling constant between the force and voltage and are used to calculate the energy landscape.

#### 9. Estimate of $k_{-T}$ and $k_T$ from ATPase data

From our model for ATPase data and our unwinding data we sought to estimate $k_{-T}$ and $k_T$ using a maximum likelihood approach. We use unwinding experiments in which $[ADP] = 0$ at 180 mV assisting force. In the high assisting force limit we ignore $k_{-SD}$ and therefore the connection matrix for the [ATP]-dependent step is:

$$M_{[ATP]-dep} = \begin{bmatrix} 0 & -k_D & k_{-D} \cdot [ADP] & 0 \\ 0 & k_D & -k_T \cdot [ATP] - k_{-D} \cdot [ADP] & k_{-T} \\ 0 & 0 & k_T \cdot [ATP] & -k_{-T} - k_{ST}(V) \\ 0 & 0 & 0 & k_{ST}(V) \end{bmatrix}. \tag{s27}$$

This model has 5 free parameters ($k_D$, $k_{-D}$, $k_T$, $k_{-T}$ and $k_{ST}$). From section s2 we know the values of $V$, $K$ and $d$. These are related to the rate constants by the following expressions (Craig *et al.* 2017 PNAS).

$$K = \frac{k_{-T} + k_{ST}}{k_T} \cdot \frac{k_D}{k_D + k_{ST}}. \tag{s28}$$

$$V = \frac{k_{ST} + k_D}{k_D \cdot k_{ST}} \tag{s29}$$

$$d = \frac{K \cdot k_{-D}}{k_D}. \tag{s30}$$

These expressions provide 3 constraint equations on the values that the rate constants can take. In the likelihood approach this leads to a reduction in dimension of the parameter space from 5 to 2. We can also use our estimates of $k_{ST}$ and $k_D$ from section s8 to further reduce the dimension of the parameter space from 2 to 1 (note: $k_{ST}$ and $k_D$ only reduce the dimension by 1 because they are related to each other and $V$ through equation s29; if we know any two of those parameters then we know the third). We therefore write the likelihood function in the single variable $k_{-T}$ as:

$$\mathcal{L}(k_{-T}|data) = \Sigma_{experiments} log(k_{ST} \cdot p_{H \cdot ATP}(t)). \tag{s31}$$

where 'data' is take from the first 6 experiments labeled dsDNA $f_{assist}$ in table s2. We calculate the log-likelihood on a lattice of guess values of $k_{-T}$ to find the maximum likelihood solution. Figure s13 shows the likelihood function and the best fit curves to the data. The likelihood function does not decay at high values of $k_{-T}$, reflecting an insensitivity of the distribution function to the value of $k_{-T}$. This can be seen in the histograms shown in figure s13: The low-likelihood solution is fairly similar to the high-likelihood solution, suggesting that the distribution functions are characterized by primarily by the values of $V$,$K$ and $d$, with $k_{-T}$ having relatively little effect, except at short dwell times and low $[ATP]$, which is also affected by the loss function. Similar phenomena were also reported in Hel308 (Craig *et al.* 2017 PNAS).

Therefore we do not report a value for $k_{-T}$ but rather use the estimate that $k_{-T} \gg k_{ST}$ to rewrite equation s28 in terms of the dissociation constant for ATP:

$$K = K_T \cdot \frac{k_D}{k_D + k_{ST}} \tag{s32}$$

which yields $K_T = \frac{k_D + k_{ST}}{k_D} \cdot K \approx 100 \pm 18\ \mu M$.

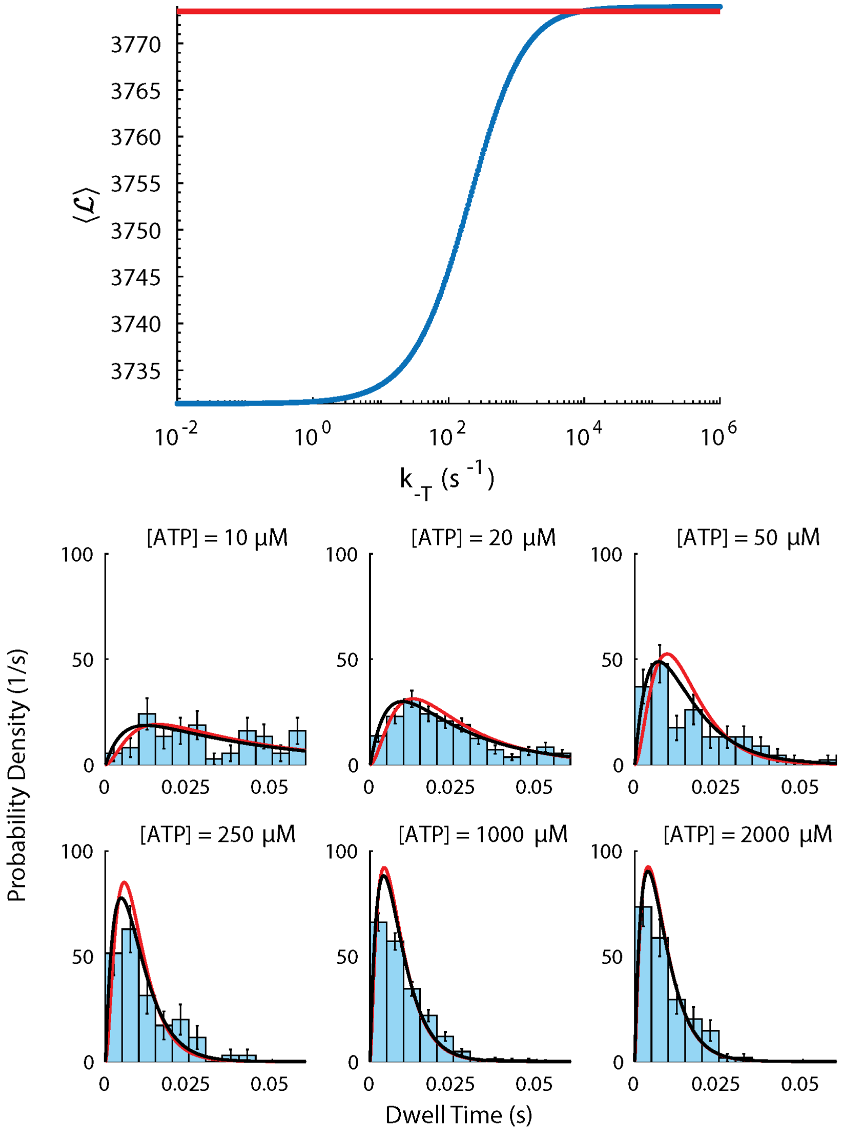


FIGURE S13. (Top) Likelihood as a function of the guess value of $k_{-T}$. (Bottom) dwell time histograms for ATP titration experiments. The red curves correspond to $k_{-T} = 10^{-4}\ s^{-1}$ (a low-likelihood solution) and the black curves to $k_{-T} = 10^{4}\ s^{-1}$ (a high-likelihood solution).

#### 10. Calculating the coordinates of the transition states and $\Delta G$ from kinetic parameters. Constructing the Energy Diagram

Figure 4 shows the energetic landscape for conformational changes of ATP-bound and ADP-bound RecQ. The physical coordinate of the transition state $x_t$ is given by the relative values of $\alpha_{ST}$, $\alpha_{-ST}$, $\alpha_{SD}$ and $\alpha_{-SD}$ and the measured step size:

$$x_{t,ATP} = \frac{\alpha_{ST}}{\alpha_{ST} + \alpha_{-ST}} \cdot x_{step,ATP} = \frac{10^{-5}}{10^{-5} + 0.014} \cdot (0.55\ nt) \approx 0.001\ nt. \tag{s33}$$

and

$$x_{t,ADP} = \frac{\alpha_{SD}}{\alpha_{SD} + \alpha_{-SD}} \cdot x_{step,ADP} = \frac{0.003}{0.003 + 0.014} \cdot (0.45\ nt) \approx 0.1\ nt \tag{s34}$$

The Gibbs energies for the conformational changes are calculated from the equilibrium constants evaluated at V = 0 mV:

$$\Delta\ G_{ATP} = -kT \cdot log(k_{ST}/k_{-ST}) = -kT \cdot log(350/11) \approx -3.5 \cdot kT \tag{s35}$$

and

$$\Delta\ G_{ADP} = -kT \cdot log(k_{SD}/k_{-SD}) = -kT \cdot log(700/2.3) \approx -5.7 \cdot kT. \tag{s36}$$

To construct figure 4B and 4D we used the values for the transition state and $\Delta G$ from above using a Piecewise Cubic Hermite Interpolating Polynomial (PCHIP). The energy minima for the ATP-bound transition are set at

$$x_{0,ATP} = 0\ nt$$

and

$$x_{1,ATP} = x_{0,ATP} + x_{step,ATP} = 0.55\ nt.$$

For the ADP bound transition

$$x_{0,ADP} = 0.55\ nt$$

and

$$x_{1,ADP} = x_{0,ADP} + x_{step,ADP} = 1.00\ nt.$$

The maxima are located at

$$x_{max,i} = x_{0,i} + x_{t,i}$$

with maximum energy arbitrarily set at $E_{max,i} = E_{0,i} + 2 \cdot kT$ for display purposes.

#### 11. Exploring how the transition state coordinate affects RecQ kinetics

To explore the effects of transition state coordinate on RecQ kinetics, we used the rate constants from table s5 but varied the location of the transition state coordinates and used equations s22-24 to determine what the response of the observables to force would be with the altered transition state. We co-varied the location of the two transition state coordinates from $x_{T,i} = 0$ to $x_{T,i} = x_{step,i}$ to explore the range of behaviors that can occur from the given rate constants.

When $x_{T,i} = 0$ there is substantially less mechanical response of the total velocity to the force, with dwell times nearly independent of force as we observe. In contrast, when $x_{T,i} = x_{step,i}$, the probability of a backwards step is nearly independent of the force while the dwell time responds much more strongly to opposing forces.

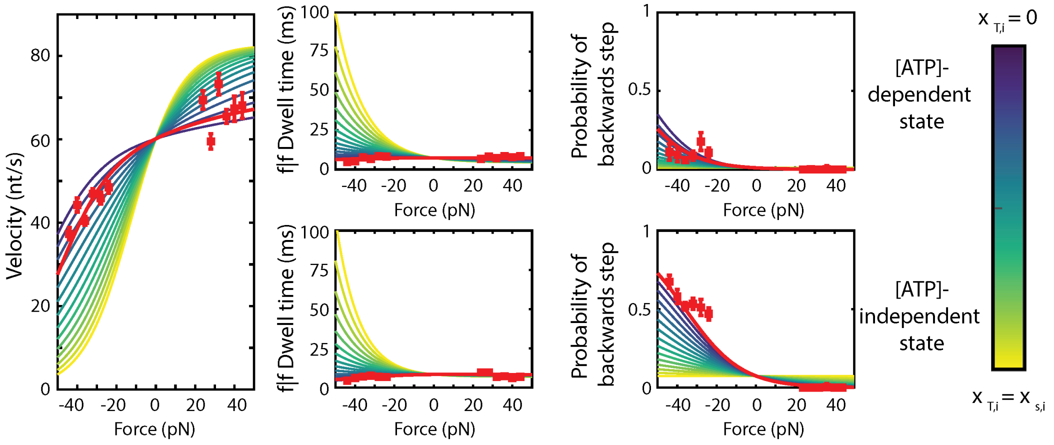


Figure s14. Velocity (left) f|f dwell times (middle) and probability of a backwards step (right) as a function of the applied force while changing the transition state. The red line is the best-fit to the data. Blue lines correspond to both transition state coordinates being located close to the pre-translocated state ($x_{T,i} = 0$). Yellow lines correspond to both transition state coordinates being located close to the post-translocated state ($x_{T,i} = x_{step,i}$).
